# Multi-site Assessment of Methods for Cell Preservation Upstream of Single Cell RNA Sequencing

**DOI:** 10.1101/2025.10.24.684427

**Authors:** Fred W. Kolling, Jessica W. Podnar, Owen Wilkins, Claire J. Fraser, Madolyn L. MacDonald, Shawn W. Polson, Zachary T. Herbert, Sridar V. Chittur, Andrew Hayden, Marcy Kuentzel, Michael Heinz, Gabriella M. Huerta, Holly S. Stevenson, Aditi Karmakar, Catrina Fronick, Lisa Cook, Sean Vargas, Xiaoling Xuei, Patrick McGuire, Molly Zeller, Yanping Zhang, Ru Dai, Xinkun Wang, Ching Man Wai, Jyothi Thimmapuram, Devender Arora, Tania Mesa, Jun Fan, Yuriy O. Alekseyev, Francis Cervone, Christopher Williams, Nickolas Gorham, Alexander Lemenze, Sara Goodwin, Jonathan Preall, Charles A. Whittaker

## Abstract

Single cell RNA sequencing (scRNA-seq) is a revolutionary technique to identify cell types and their molecular phenotype in heterogeneous biological specimens. ScRNA-seq typically requires fresh, high quality single cell suspensions that are processed immediately to preserve their molecular profiles. This presents a challenge for samples with long preparation times and prevents collection at remote sites lacking the required instrumentation for sample processing. Recently, several commercial assays have been released that enable sample preservation at the time of collection either via fixation or cryopreservation, allowing for sample processing to occur months after the initial collection. The Association of Biomolecular Research Facilities’ (ABRF) DNA Sequencing (DSRG) and Genomics Bioinformatics (GBiRG) Research Groups have undertaken a cross-platform, multi-site study to assess the performance and reproducibility of three platforms: a) 10x Genomics FLEX, b) Parse Bioscience Evercode WT v2 and c) Honeycomb Bio HIVE. Total leukocytes were isolated from a single healthy individual using the EasySep RBC depletion reagent. Cells were then characterized by collecting a 21-color flow cytometry dataset for reference and the remaining material was used for scRNA-seq procedures where different sites then processed either the fixed or cryopreserved cells for each method. We evaluated performance of each method across traditional scRNA-seq quality control metrics and analysis applications, including gene/transcript detection sensitivity, cell type discovery and annotation, and differential expression. We demonstrate that data from the methods tested can be effectively integrated and produce concordant results with regard to cell type annotation and relative abundance, though we observe platform-specific differences in the expression of a subset of genes. Preservation-based methods also show better retention of fragile granulocyte populations compared with fresh samples processed using the 10x 3’ workflow. The improvements to preservation methods are changing the way research is conducted and our thorough investigation into the performance of each method provides a valuable resource to help scientists determine the most appropriate single cell preservation workflow given their sample collection logistics and laboratory infrastructure constraints.

## Introduction

In recent years, single cell RNA sequencing (scRNA-seq) has emerged as a powerful tool to measure gene expression within individual cells, providing the resolution required to determine cell identity and define cell states. These methods rely on microfluidically-generated droplets (Matuła et al. 2020), picowells (Gierahn et al. 2017), or cells themselves (Wohnhaas et al. 2019), to partition and uniquely barcode the RNA molecules within each cell (Rosenberg et al. 2018; Wohnhaas et al. 2019). The barcoded RNAs are then sequenced to determine their identity and the cell of origin. The starting material for these experiments is a suspension of single cells, produced either through the dissociation of tissues, or purification of cells from biofluids (e.g. blood). The integrity of the cells in suspension is critical, and has a direct impact on the quality of data from scRNA-seq experiments. Cells that are stressed or undergoing apoptosis generate distinct gene expression signatures and actively degrade their mRNAs, reducing transcriptome complexity (Wu et al. 2021; Ilicic et al. 2016; Zhao 2002). Furthermore, cell lysis causes the release of RNAs into the cell suspension and these “ambient” RNAs can become barcoded and erroneously associated with cell expression profiles, decreasing the cell-specific signal (Fleming et al. 2023; Young and Behjati 2020). For these reasons, single cell experiments have required optimized dissociation procedures using fresh, high-viability single cell suspensions for use in these assays. This presents a challenge when large numbers of samples are processed in parallel, or when the reagents and instrumentation for single cell capture are distant to where the cell suspension is generated. To circumvent these issues, methods such as methanol fixation and cryopreservation have been validated in specific sample types (e.g. blood, cerebrospinal fluid, etc.) (DuBois et al. 2024; Touil et al. 2023; Gutiérrez-Franco et al. 2023; Chen et al. 2021; Wohnhaas et al. 2019; Alles et al. 2017) however their impact on cell type representation and transcript detection in some contexts has prevented their widespread use (Denisenko et al. 2020).

Recently, several commercial solutions including 10x Genomics FLEX, Parse Bioscience Evercode, and Honeycomb Bio HIVE have come to market to enable preservation of single cell samples upstream of scRNA-seq. Separating the preparation of single cell suspensions, from cell capture, library preparation and sequencing, makes it possible to collect samples in one location and ship to another for processing. As a result, core facilities and other service providers will be eager to implement these methods to simplify sample collection logistics for internal users and to facilitate access by external clients to single cell services. However, there are many variables to consider when choosing a platform including sample preservation (fixation or cryopreservation) and RNA capture (Poly-dT priming, random priming or probe-based detection) methods between platforms, as well as the instrumentation and hands-on time required. Differences in cell capture and sequencing performance have been described in comparative studies evaluating various scRNA-seq methods (De Simone et al. 2024; Xie et al. 2024; Hornung et al. 2023; Ding et al. 2020). This study provides additional insight into the impacts of prolonged storage time and kit reproducibility across multiple performance sites, as well as considerations for selecting a given chemistry.

To evaluate the performance, reproducibility and workflow requirements of the available methods, the Association for Biomolecular Research Facilities’ (ABRF) DNA Sequencing (GSRG) and Genomics and Bioinformatics (GBiRG) Research Groups have conducted a multi-site, cross-platform assessment of the 10x Genomics FLEX, Honeycomb HIVE and Parse Evercode technologies. We show that while there is general agreement in cell type assignment, relative abundance and overall gene expression profiles across platforms and performance sites, strong technology-specific expression signatures exist within the data and cross-site reproducibility varies between methods. In addition, we find that sample preservation by fixation (FLEX) and freezing (HIVE) improves the retention of fragile granulocyte populations compared with fresh sample processing using the traditional 10x 3’ GEX (3pGEX) assay. This study demonstrates the importance of selecting an appropriate single cell technology based on experimental and logistical constraints, and provides a resource to core facilities and individual users to make informed decisions when implementing these assays.

## Methods

### Study Design

The study was designed to evaluate reproducibility, ease of use and performance of three technologies for cell preservation on the same sample type across three commercially available options. We selected a single site to collect and preserve the samples, and nine ABRF-member genomics core facilities to perform downstream processing. To assess variability in sample preservation, preservation protocols were performed by two different technicians in parallel to produce “A” and “B” replicates. Once preservation protocols were completed the samples were stored at -80 C for two weeks before shipping to each of the participating sites. Prior to any site receiving their samples, they were provided with hands-on training by the respective vendors using control samples. After two weeks, each site received “A” and “B” preserved samples and stored for an additional two weeks at -80 C, for a total of 4 weeks storage time before completing the assigned workflow. Sequencing ready libraries were shipped to a central site for quality assessment and sequencing. Our follow-on study using PBMCs was performed at a single site following the same protocols described for the total leukocyte experiment except where noted below.

### Sample Collection, Cell Isolation and QC

Blood collection from a single healthy male was performed at Dartmouth Hitchcock Medical Center under an IRB approved protocol. Approximately 50 ml of whole blood was collected into EDTA vacutainer tubes (BD Bioscience) and immediately transported to the Immune Monitoring Lab at Dartmouth Cancer Center for processing. For total leukocyte isolation, the blood sample was subjected to two rounds of RBC-depletion using magnetic beads using the EasySep RBC Depletion kit (Stem Cell Technologies) generating about 8 x 10^^7^ cells. Two additional washes were performed with samples spun at 500 x g for 10 minutes to reduce the presence of platelets. For PBMC isolation, gradient separation of the blood was performed using Histopaque-1077 (Corning). Briefly, blood was overlaid onto the Histopaque-1077, then centrifuged for 30 minutes at 750 x g. An additional spin performed with samples at 500 x g for 10 minutes to reduce the presence of platelets. For both sample types, cells were transferred to the Dartmouth Genomics Shared Resource for counting and viability assessment on a Nexcellom K2 instrument (Revvity), prior to running each of the workflows described below. Both total leukocyte and PBMC samples exhibited >95% viability as determined by acridine orange/propidium iodide (AO/PI) staining.

### Flow Cytometry

Cells were stained for flow cytometry analysis as follows. 2×10^6 total leukocytes or PBMC were suspended in 1.25 µg/ml of human IgG to block Fc receptors and incubated for 10 minutes in a 5 ml FACS tube. Following this, 5µl of Brilliant Stain Buffer Plus (BD Biosciences) and 5µl of True-stain Monocyte Blocker (Biolegend) were added in succession. Due to steric hindrance preventing it from being used in a master mix of antibodies, anti-TCRγδ was added, and cells incubated for 10 min at RT in the dark. The remaining antibodies were then added in a master mix and the cells incubated for 30 min at RT in the dark. All antibodies used are listed in Supplemental Table 1. To wash unbound antibodies, 3 ml of PBS was added and cells centrifuged at 500 x g for 5 minutes. Cells were resuspended in 1% paraformaldehyde in PBS to fix antibodies to cells and then washed as above. Cells were then acquired on a 5-laser Aurora Spectral Cytometer (Cytek).

### 10x Genomics 3’ v3.1 on Fresh Specimens

Following QC, cells were loaded onto two separate lanes of a Single Cell Chip G to generate “A” and “B” replicates, and processed on a 10x Chromium X instrument, targeting 10,000 cells per sample. Emulsions containing encapsulated single cells were removed from the chip and processed according to the Chromium NextGEM Single Cell 3’ v3.1 User Guide (CG000204 RevD). Completed libraries were examined on a Bioanalyzer (Agilent) and quantified by Qubit (Thermo Fisher) prior to pooling and loading on a Novaseq6000 instrument as described below.

### Sample Preservation

Following collection, cell isolation and QC, samples were processed with their respective preservation protocols - cell fixation or cryopreservation v- and stored at -80C prior to scRNASeq library generation. To ensure the highest cell quality and minimize sample variation for all methods, a plan was formulated to process the samples for preservation in parallel at the central site followed by distribution to the testing sites. Details can be found in the Cell Preservation Supplemental Protocol (Fred Kolling IV 2024). Cells for the 10x FLEX RNA profiling kit followed the Fixation of Cells & Nuclei for Chromium Fixed RNA Profiling demonstrated protocol (CG000478, Rev A) using 1 million cells per fixation. Cells for the Parse workflow were fixed using the Evercode Cell Fixation v2 kit (ECF2001) following the Evercode Fixation user manual, V2.0.1 with 3 million cells per fixation. Honeycomb HIVEs were loaded with 15K cells into v1 HIVEs following the HIVE scRNASeq v1 sample capture user protocol (Rev A) for the leukocyte samples, and 30K into the CLX Hives for PBMCs following HIVE CLX scRNAseq Sample Capture protocol.

### 10x Genomics FLEX Sample Processing on Fixed Cells

10x FLEX single cell libraries were completed with the Chromium Fixed RNA Profiling kit, part number 1000474, to process fixed cells following user guide, CG000477 Rev A. Cell fixations were completed by a central site and aliquots with 1 million fixed cells shipped on dry ice to each of the testing sites. Briefly, fixed and permeabilized cells underwent probe hybridization and ligation for 16 hours, partition of Gel Beads-in-emulsion (GEM) and barcoding, cDNA amplification, and library construction each with unique indices following the protocol of 10X Chromium Fixed RNA Profiling Reagent Kits User Guide, CG000477 RevA (10X Genomics). For the leukocytes, each testing site processed replicate “A” and “B” to produce a total of 8 libraries and a single library was generated for the PBMC sample. The targeted recovery per sample was 10K cells per library. The quality and quantity of each library was checked with the Qubit (ThermoFisher Scientific), Agilent Bioanalyzer 2100 HS DNA kit (Agilent Biotechniques) and qPCR using the Kapa Sybr Fast RT-qPCR kit, KK4602 (Roche Diagnostics).

### Honeycomb Hive Sample Processing

Honeycomb single cell libraries were completed following the HIVE scRNAseq V1 or CLX Processing kit user protocols (Honeycomb Biotechnologies, Inc.). V1 kit Rev A was used for leukocytes and CLX for the extension study with the PBMCs. The central site that loaded the HIVEs used the HIVE scRNASeq complete kit (HCB018) that included reagents and materials for sample loading and processing while the testing sites used the HIVE scRNAseq Starter Bundle (HCB019) for sample processing only. HIVEs were shipped from the central site to testing sites on dry ice for processing. Briefly, on day 1 the cell loaded HIVE Collectors were thawed and washed followed by cell lysis and a hybridization step to capture Poly-A transcripts from individual cells in the HIVE picowells on the beads. The beads were recovered following the HIVE protocol and then transferred to a 96 well filter plate to perform 1st and 2nd strand synthesis. Steps were completed in the 96 well filter plate utilizing vacuum assembly provided by the manufacturer as part of the Hive scRNASeq Starter Bundle. Following 2nd strand synthesis a whole transcriptome amplification step was performed. Protocol was continued on day 2 to complete the library prep, SPRI clean up of WTA product followed by an additional PCR to incorporate the Illumina sequencing adapters and indices using a total of 25ng per reaction. For the leukocytes, each testing site processed replicate “A” and “B” to produce a total of 8 libraries and a single library was generated for the PBMC sample. The quality and quantity of each library was checked with the Qubit (ThermoFisher Scientific), Agilent Bioanalyzer 2100 HS DNA (Agilent Biotechniques) and RT-qPCR using the Kapa SYBR Fast qPCR kit, KK4602 (Roche Diagnostics).

### Parse Biosciences Sample Processing

Parse single cell library was prepared with the Parse Single Cell Whole Transcriptome kit, Evercode WT Mini v2, ECW02010 (Parse Biosciences) following the user manual V2.0.1. For the first round of barcoding, 4,000 cells were distributed into each of 12 wells (48,000 cells total) with subsequent pooling and barcoding steps performed following the user guide. Two sub-libraries were completed for the sample and a single sub-library was sequenced. The quality and quantity of the sequencing library was checked with the Qubit (ThermoFisher Scientific), Agilent Bioanalyzer 2100 HS DNA kit (Agilent Biotechniques).

### Illumina Sequencing

Sequencing was completed at one site using the NovaSeq 6000 instrument from Illumina. Libraries generated from the leukocytes for 10x were sequenced on a S2 100 cycle flow cell 28-10-10-90 targeting 10K reads per cell for FLEX and 25K reads per cell for the 3’ GEX libraries. Pooled loading concentration for 10x libraries standard workflow was 1500 pM. The Honeycomb leukocyte libraries were sequenced on a full S1 100 cycle flow cell 25-8-8-5 targeting 25K reads per cell. Pooled loading concentration for Honeycomb libraries standard workflow was 1800 pM. 10x and Parse PBMC libraries were sequenced on individual lanes using the XP workflow with a SP 200 cycle flow cell 100-10-10-100, pooled loading concentration was 750 pM and the Honeycomb PBMC library sequenced on a full SP 100 cycle flow cell, 25-8-8-50. Prior to sequencing, qPCR was completed to determine library concentration for pooling prior to loading the sequencer. The Honeycomb libraries required custom primers which are provided in the sample processing kit with the library prep reagents at 100 uM. The NovaSeq 6000 Custom Primer protocol, Document 1000000022266 v03, was followed to prepare and load the custom primers for the Honeycomb sequencing runs.

### Data Analysis

#### Single cell RNA-seq preprocessing

Sequencing data (FASTQ format) was processed to generate cell-level feature barcode matrices using vendor software. For leukocyte samples, the 10x samples were processed using cell ranger v7.1.0 and data from all valid barcodes (raw_feature_bc_matrix data) was used in downstream analysis. Honeycomb leukocyte data was processed using BeeNet v1.1.3 specifying 10,000 barcodes. The 10x PBMC samples were processed using Cell Ranger v7.1.0 and only cell-associated barcodes were used for downstream analysis (filtered_feature_bc_matrix data). The PBMC Honeycomb sample was processed using BeeNet v1.1.3 specifying 15,000 barcodes. The PBMC Parse Evercode data was processed using split-pipe v1.0.5p with default settings.

Following data generation, processing and analysis were carried out in a containerized R environment that includes a variety of data science, genomics and single-cell RNA-Seq software packages (https://github.com/GBIRG/scRNAseq_2022/tree/main/Docker, Fig. S1). The containerized compute solution we have implemented facilitates distributed and collaborative data analysis projects and the image is made available to facilitate public analysis of this dataset (docker://alemenze/abrfseurat). A listing of all the packages and versions is available in the supplementary file DSRG_GBIRG_Rcontainer_sessionInfo.txt. In order to capture low-information content granulocytes in the leukocyte samples, data were imported into ‘Seurat’ version 4.3.0 (Hao et al. 2021) using the following low-stringency parameters:nFeature_RNA >= 30, nCount_RNA >= 100, mitoRatio >= 0.1, and genes with non-zero counts in at least 10 cells. The data from each platform were then normalized using *sctransform* (Hafemeister and Satija 2019) and integrated using anchor-based integration (Stuart et al. 2019) followed by dimensionality reduction and clustering.

#### Cell type assignment

Clusters were then characterized with an integrated assessment of cluster-specific marker genes, transcriptional complexity, and scoring of expression signatures with ‘singleR’ (Dvir Aran, Aaron Lun, Daniel Bunis, Jared Andrews, Friederike Dündar, n.d.). In order to assign cell types to each leukocyte (Fig. 3A) and PBMC (Fig. 3C) cluster, ‘singleR’ was used with the ‘celldex’ (Aran et al. 2019) references: HumanPrimaryCellAtlas (Mabbott et al. 2013), DatabaseImmuneCellExpression (Schmiedel et al. 2018), and BlueprintEncode (Martens and Stunnenberg 2013). ‘SingleR’ assigns identity at the cellular level. To assign cell types to unsupervised clusters, the fraction of cells assigned to each identity was calculated for each cluster and assignments that were coherent on a cluster-level were prioritized. In addition to ‘singleR’, cluster-specific gene markers were identified using *FindAllMarkers* and these marker lists were examined using the MsigDB Investigate Gene Sets (Wilson et al. 2014; Liberzon et al. 2015) utility and the Cell Type Signature (C8) collection. Cell type assignment conclusions were then manually validated by examination of cell type specific markers. Debris (cell-free or empty droplets) clusters were defined as those dominated by cells or droplets with poor QC metrics, ambiguous cell type assignments using ‘singleR’ and no cluster-specific markers. Clusters meeting these criteria were flagged as debris and excluded from downstream analyses (Fig. S2).

To remove potential sequencing depth-related biases in downstream analysis, the sample-level cell counts following debris exclusion were used as denominators to downsample the raw data to a sequencing depth of 25,000 reads per cell. The resulting FASTQ files were then used as input to the same analysis procedure. For the PBMC data, vendor software was used to exclude debris clusters from the 10x 3pGEX and FLEX, and Parse samples. For the HIVE samples, 15,000 cells were requested in processing and a similar cluster-level debris exclusion process as used in the leukocytes was applied. As in the case of the leukocytes, once a reliable cell count was obtained, FASTQ files were downsampled to 25,000 reads per cell and analysis was repeated. Cell type assignments prepared for the pre-downsampling object were then transferred to the downsampled object so that cells flagged as low-confidence coils be excluded from the analysis of downsampled data. These metadata were then re-evaluated in the context of the downsampled data to revise and finalize cell type assignments.

For comparison of transcript coverage over gene bodies, the Python package deeptools v3.5.4 (Ramírez et al. 2016) was used to generate BigWig coverage files from the BAM-formatted alignments for each sequencing platform. Normalized coverage over exons was computed using deeptools *computeMatrix* with the parameters: scale-regions -b 100 -a 100 -- unscaled5prime 60 --unscaled3prime 60 --skipZeros –metagene and gene annotations GTF file provided by 10X Genomics (GRCh38-2020-A, based on GENCODE v32/Ensembl 98).

#### Correlation analysis

To agnostically evaluate the similarity between samples across technologies and sites, we applied a bulk correlation analysis across the dataset. SCT normalized expression counts were aggregated per-sample using the ‘Seurat’ function *AggregateExpression* on SCT assay data and Pearson correlations were calculated between each pair-wise sample for downstream visualization.

#### Differential expression analyses

To perform multi-site, cross-platform assessment, we executed comparisons between A and B replicates generated at each site, between A replicates from each site and 28 day and 1 day storage time. To ensure consistent comparisons across analyses, we filtered to only genes present in the FLEX-probeset, which includes 28,690 genes to make the comparisons uniform across each platform. For each comparison, count data for relevant replicates were extracted from the object and differential expression analysis using ‘DESeq2’ (Love et al. 2014) and a∼condition design was then performed. Differentially expressed genes (DEGs) were defined as those having an absolute log2 fold change (log2FC) > 1 and an FDR adjusted p-value (padj) < 0.05. The results we visualized using enhanced volcano plots were generated for each comparison using ‘EnhancedVolcanò (Blighe 2018) and fold change box plots showing A_vs_B and Site-specific variation were generated using ‘ggplot2’ (Wickham 2016).

#### UMI gene body coverage analysis

For sequencing saturation analyses, BAM files were downsampled to a range of different depths relative to the starting library using Pysam v0.18.0. Reads were then re-counted for unique UMI/gene/barcode sets. In brief, only reads originally tagged by Cell Ranger with the ‘xf’ tag equal to 17 (read mapped confidently to the transcriptome and to 1 unique feature) or 25 (read mapped confidently to the transcriptome and to 1 unique feature and read is representative for a molecule and is treated as a UMI count) were added to sets for each gene/barcode pair, resulting in a matrix that faithfully reproduces the Cell Ranger counting rubric at each depth. To extrapolate the number of predicted mean UMIs/cell at saturation, downsampled data was fit to a Michaelis-Menten model of the form:

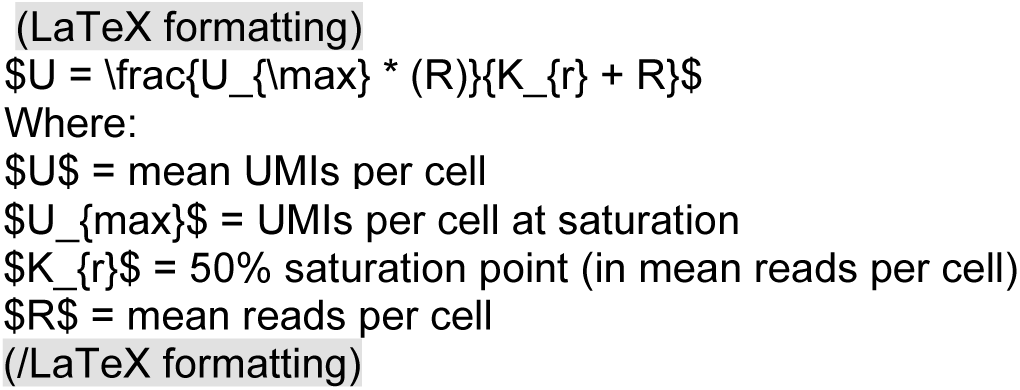

## Results

We compared data generated by three different single-cell RNA-Seq technologies (10x FLEX, Honeycomb HIVE and Parse Evercode) that enable users to preserve cells prior to library preparation. Total human leukocytes (PBMCs + granulocytes) were profiled due to the well-established cell type-specific expression profiles of PBMCs, and the potential for preservation focused scRNA-seq protocols to capture granulocytes, which have been underrepresented in single-cell data to date due to their fragility and low transcriptional complexity. To establish a reference to many existing datasets, fresh leukocytes were processed using the 10x Genomics 3’ v3.1 chemistry (3pGEX). To minimize the number of variables in our study, the total leukocyte and PBMC samples were collected from the same individual at a single site. Fixation and storage were performed in parallel by two technicians (designated “A” and “B” replicate samples) to assess technical variability in sample prep. Preserved samples from the FLEX and HIVE were shipped to 4 separate sites and processed after 28 days in storage to assess the reproducibility of these methods and identify possible sources of cross-site variation. Kit compatibility challenges with total leukocytes in the Parse Evercode chemistry prevented their inclusion in the initial phase of this study, so a second phase was conducted to generate data from all 3 platforms. This follow on experiment was performed on PBMC specimens isolated at the same site as the first phase and processed using the 10x 3pGEX (fresh), 10x FLEX, HIVE and Evercode workflows. Table 1 presents the experimental metadata associated with the 20 leukocyte samples and 4 PBMC samples. Each sample was sequenced at a central location to the manufacturer recommended depth (Table 1, ReadCount, Fig. S3A). Count data were prepared using vendor-specific tools and the resulting data were imported into ‘Seurat’ v4 (Hao et al. 2021) for analysis.

**Table 1.**
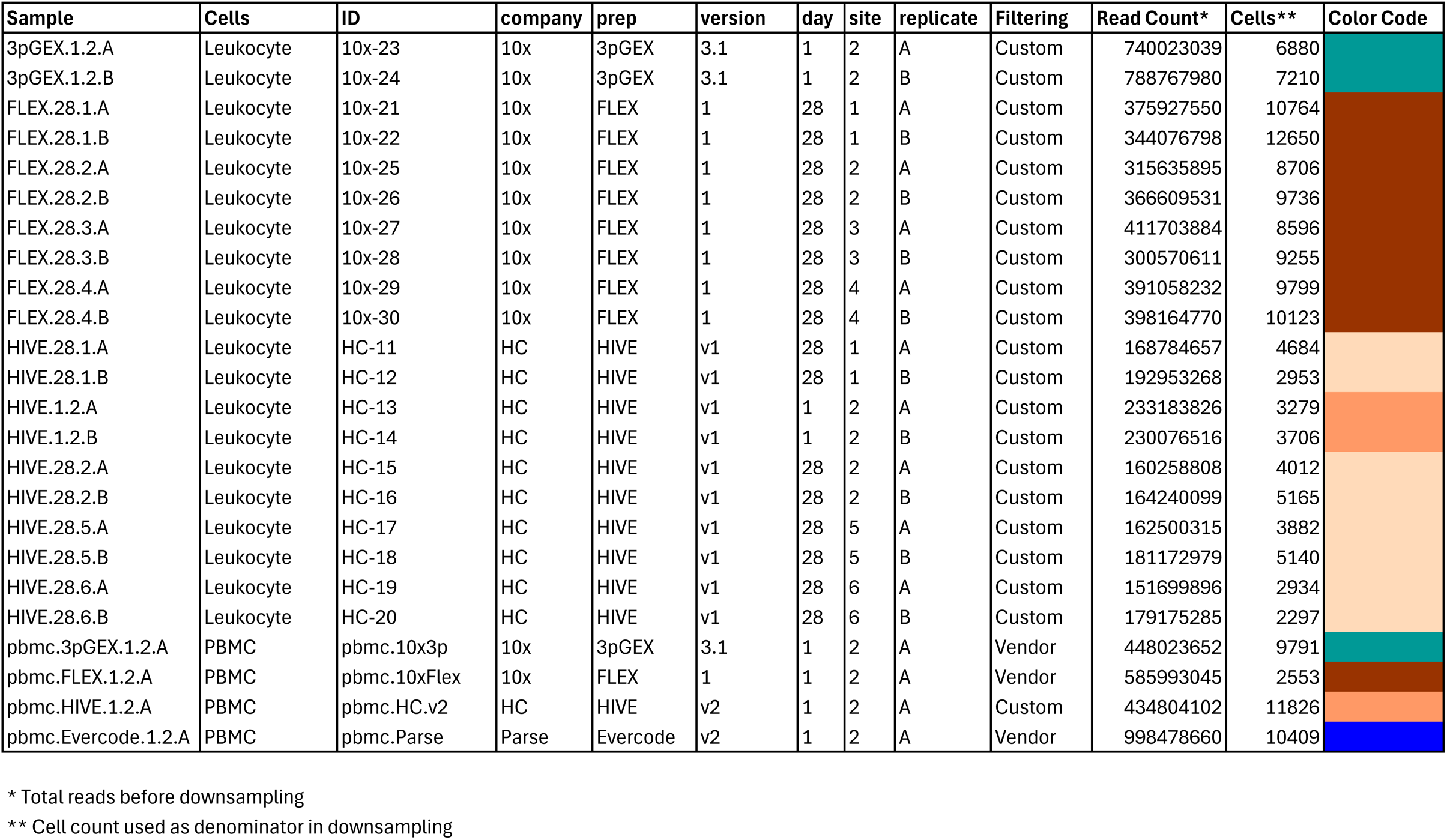
Experimental Metadata. A total of 24 samples were considered in this experiment, 20 leukocyte samples and 4 PBMC samples. The nomenclature presented in the Sample column is used in sample-level visualizations. The color key is used to indicate experimental groups in subsequent figures.

With the exception of the 10x and Parse PBMC samples, the initial processing used permissive filters for transcriptional complexity to ensure retention of granulocytes. These unfiltered data were then integrated, clustered, and cluster-specific markers were identified by differential expression analysis. Cell-type assignment was performed using ‘singleR’ and by examination of cluster-specific marker genes. Clusters with poor QC parameters, ambiguous cell types and without clear distinguishing marker genes were flagged as cell-free droplets and excluded from downstream analysis (Fig. S2). To standardize the per-cell sequencing depth for different samples (Fig. S3C), the counts of high confidence cells (Table 1, Cells, Fig. S3B) were then used as a denominator to downsample the original FASTQ files to a consistent sequencing depth of 25,000 reads per cell (Fig. S3D).

Following downsampling of the FASTQ files and re-processing using vendor software, data were subset to droplets labelled as cells in the initial analysis of the full dataset. Traditional quality control metrics (number of genes per cell (n_feature), number of UMIs per cell (n_count) and the fraction of reads derived from the mitochondrial genome (mitoRatio)) were evaluated to compare each platform. For the leukocyte data (Fig. 2A), the site-level A and B replicates are highly consistent with the exception of those from site 4 (FLEX.28.4A and FLEX.28.4B, red arrows). In addition to this FLEX replicate variability for site 4, there is variability in both the number of genes and the number of counts within the FLEX sets. Samples from sites 1 and 4 (FLEX.28.1.A/B and FLEX.28.4.A/B) have more genes and counts than samples from sites 2 and 3 (FLEX.28.2.A/B and FLEX.28.3.A/B). The number of genes and counts observed in the site 1 and 4 FLEX samples is similar to what was observed in the 10x 3pGEX control.Transcriptional complexity was consistent across all four HIVE 28 day site samples, however higher counts were observed in 1 day samples compared to those after 28 days of storage and this difference may be pronounced in monocytes, B-cells and DCs. During import, cells with greater than 10% mitochondrial reads were excluded from consideration and that is reflected in the plots of this parameter. Notably, the 10x FLEX technology lacks probes targeting mitochondrial ribosomal protein genes which account for the majority of mitochondrial transcripts. This results in a lower proportion of reads attributed to mitochondrial expression (red arrowheads). In the case of PBMCs (Fig. 2B), FLEX had slightly more features detected than 3pGEX, with HIVE and Evercode showing lower detection. Interestingly, HIVE exhibited a wider distribution of both features and total counts. Similar sample characteristics were observed at the cell-type level for both leukocytes (Fig. S4A) and PBMCs (Fig. S4B) including consistent distribution of cell types across sites and technologies, suggesting successful integration of datasets (Fig. S5 A and B). The exception is the presence of platelets and an “unknown” cluster likely containing debris or ambient RNA in the HIVE PBMC sample (Fig. S5B), likely accounting for the wider distribution of feature counts in that dataset.

**Figure 1.**
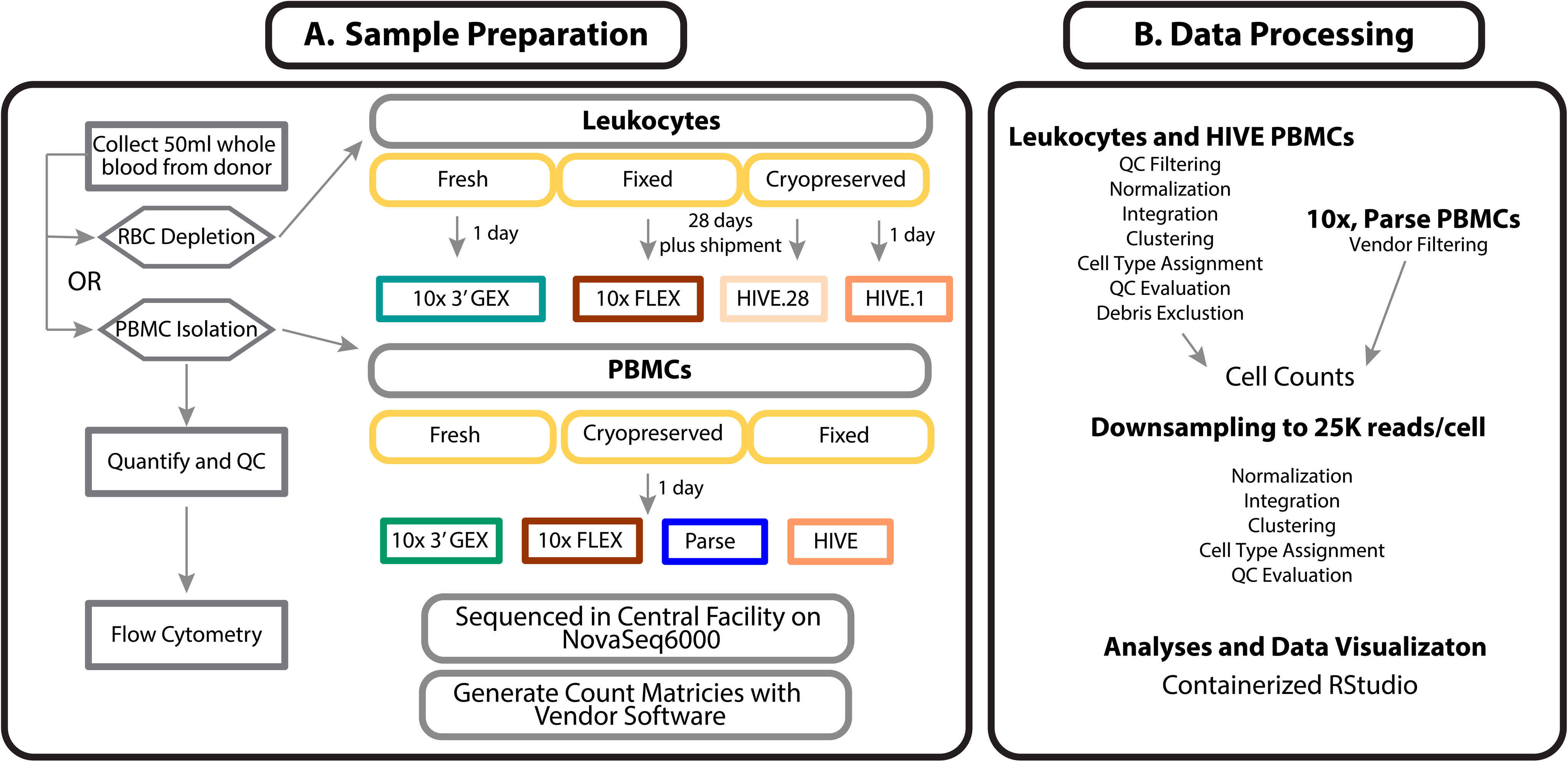
Experimental design and data processing. Leukocyte and PBMC samples obtained from a single donor were used fresh in the 10x Genomics NextGEM 3’ (3pGEX) procedure or preserved according to manufacturer workflows for the 10x FLEX, Honeycomb HIVE, and Parse Evercode (PBMC only) samples. The Leukocyte samples were shipped to four separate sites per kit, while PBMC processing was performed at a single site. Following library preparation and sequencing, Leukocyte single cell data was imported into ‘Seurat’ using minimal filtering parameters to ensure that low information-content granulocytes were not excluded by default vendor filtering. After this initial processing, clustering and cell-type assignment was performed and clusters consisting of low-quality cells with poor QC characteristics were excluded from downstream analyses. For PBMCs, a similar approach was applied to the HIVE data because the vendor software requires specification of the expected number of cells and this results in inclusion of some low-quality cells. For the 10x and Parse PBMC samples, default vendor processing was used. were processed separately by normalization, integration and clustering. Debris clusters were defined as those having poor QC characteristics.

**Figure 2.**
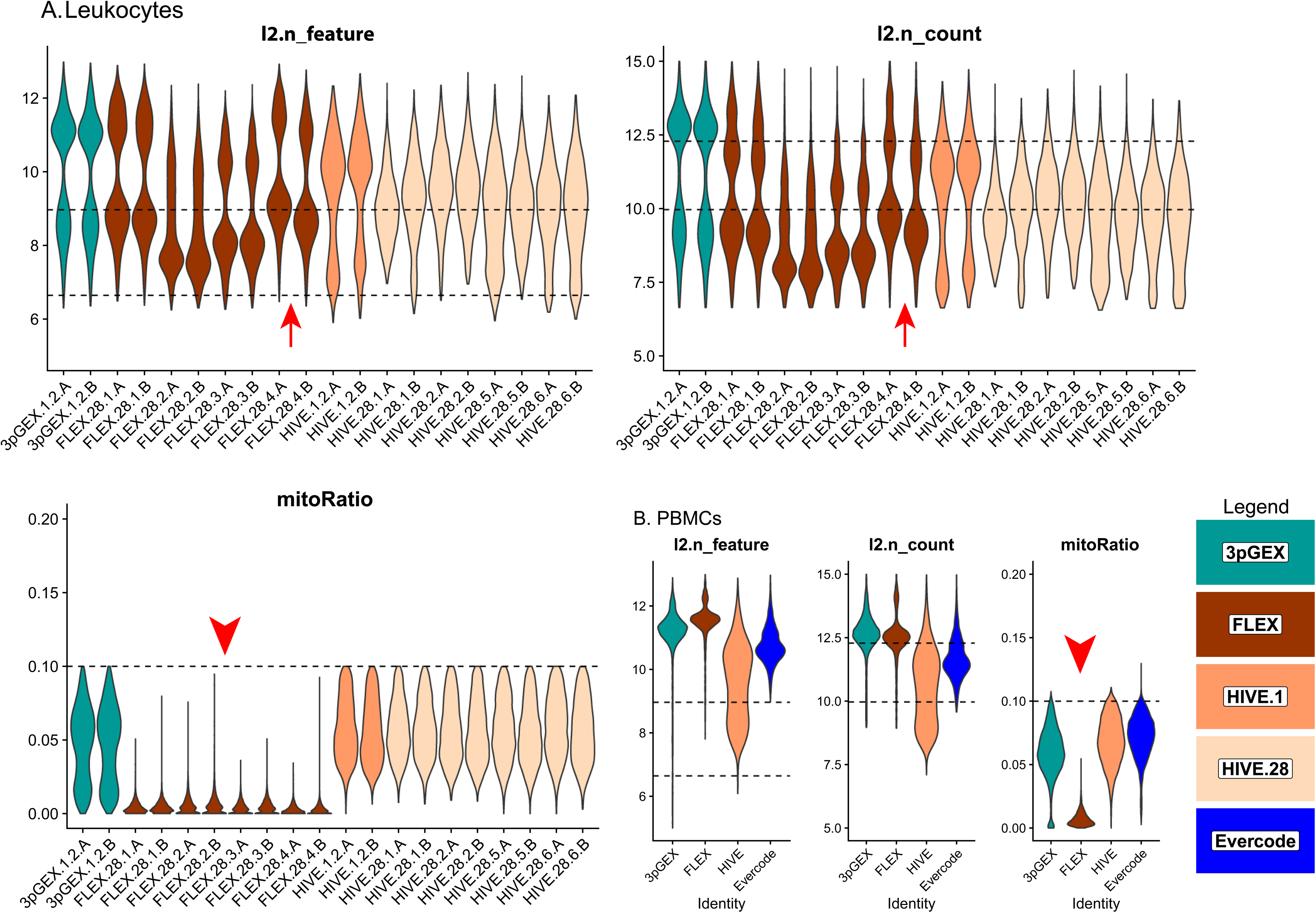
Quality control statistics for Leukocyte and PBMC samples. Violin plots presenting the log2 transformed gene count (l2.n_feature), read counts (l2.n_counts) and the percent mitochondrial reads (mitoRatio) for filtered and downsampled leukocyte (A) and PBMC (B) samples. Site-level replicates have consistent values with the exception of FLEX.28 site 4 where replicate A is slightly higher in gene and read counts compared to replicate B (Red Arrows). Mitochondrial read percentages are relatively low in the 10x FLEX samples because probes interrogating these genes are excluded from the platform (Red Arrowheads). Violins are colored according to preparation. Dashed reference lines are provided at 6.6 (100) and 9 (500) for gene counts in l2.n_feature panels, 10 (1000) and 12.3 (5000) for read counts in l2.n_count panels and 0.1 (10%) for percent mitochondrial reads in mitoRatio panels.

To facilitate comparison of cell type proportions estimated by scRNA-Seq and (flow cytometry assisted cell sorting (FACS), the detailed assignments were collapsed to generalized cell types (Fig. 3 B and D).. Overall, cell type proportions between the scRNA-Seq and FACS were generally concordant. Approximately 50% of the cells in the Leukocyte groups are granulocytes, with the majority being neutrophils. As expected, this population is a much lower fraction in the PBMC samples. In the leukocyte data, there is close agreement between the HIVE 1 day and HIVE 4 week groups indicating that this storage period does not impact cell type proportions. 10x 3pGEX demonstrated the lowest proportion of granulocytes; they are still abundant. T- and NK-cell proportions were also enriched in 10x 3pGEX samples compared to all other single-cell technologies, and more closely resembled the proportions of these cell types from FACS data. In contrast, FLEX and HIVE data demonstrated enriched proportions of monocytes relative to FACS and 10x 3pGEX. For the PBMC samples, we note strong agreement between the FACS data and both 10x technologies, while HIVE and Evercode technologies exhibited a reduction in the proportion of T-cells present and an increase in the proportion of monocytes. All single-cell technologies demonstrated a slight enrichment of B-cell proportions relative to FACSTogether, these results suggest single cell preservation platforms achieve similar cell type composition to each other and FACS data, however some exceptions exist for specific cell types.

**Figure 3.**
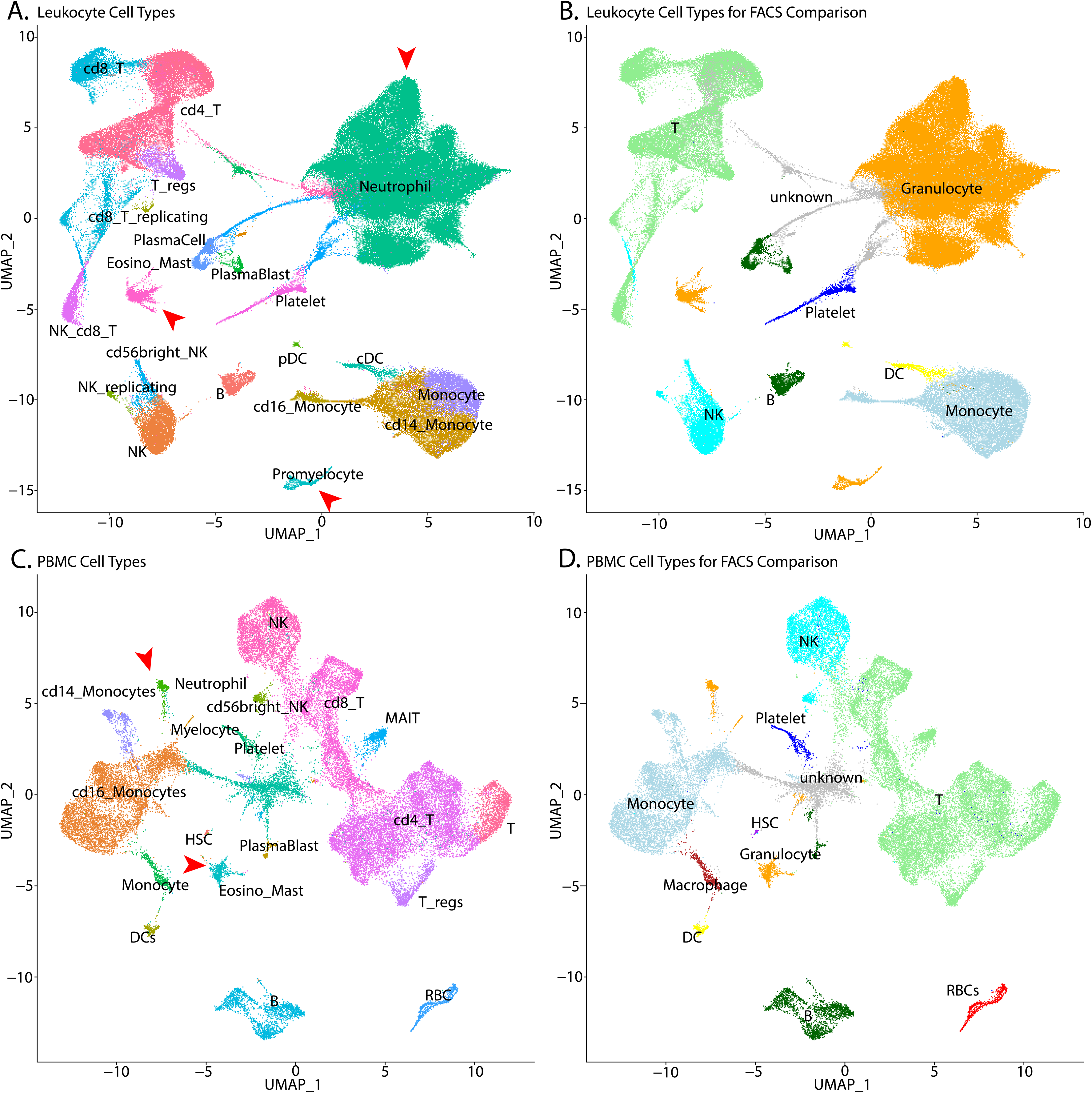
UMAP plots with cell-type assignments of clusters. UMAP plots of leukocytes (A and B) and PBMCs (C and D) with detailed cell assignments (A and C) and more general assignments used for comparison to FACS data (B and D). The color scheme used in panels A and C is the same as the color scheme used in panels B and D. The number of granulocytes (A and C, red arrowheads; B and D, orange cells), specifically neutrophils, are larger in the Leukocyte samples. T=T-cell, B=B-cell, NK=natural killer cells, DC=dendritic cells, pDC = plasmacytoid dendritic cell; cDC = conventional dendritic cell; NK = natural killer cell; HSC = hematopoietic stem cell, RBC=Red Blood cells.

**Figure 4.**
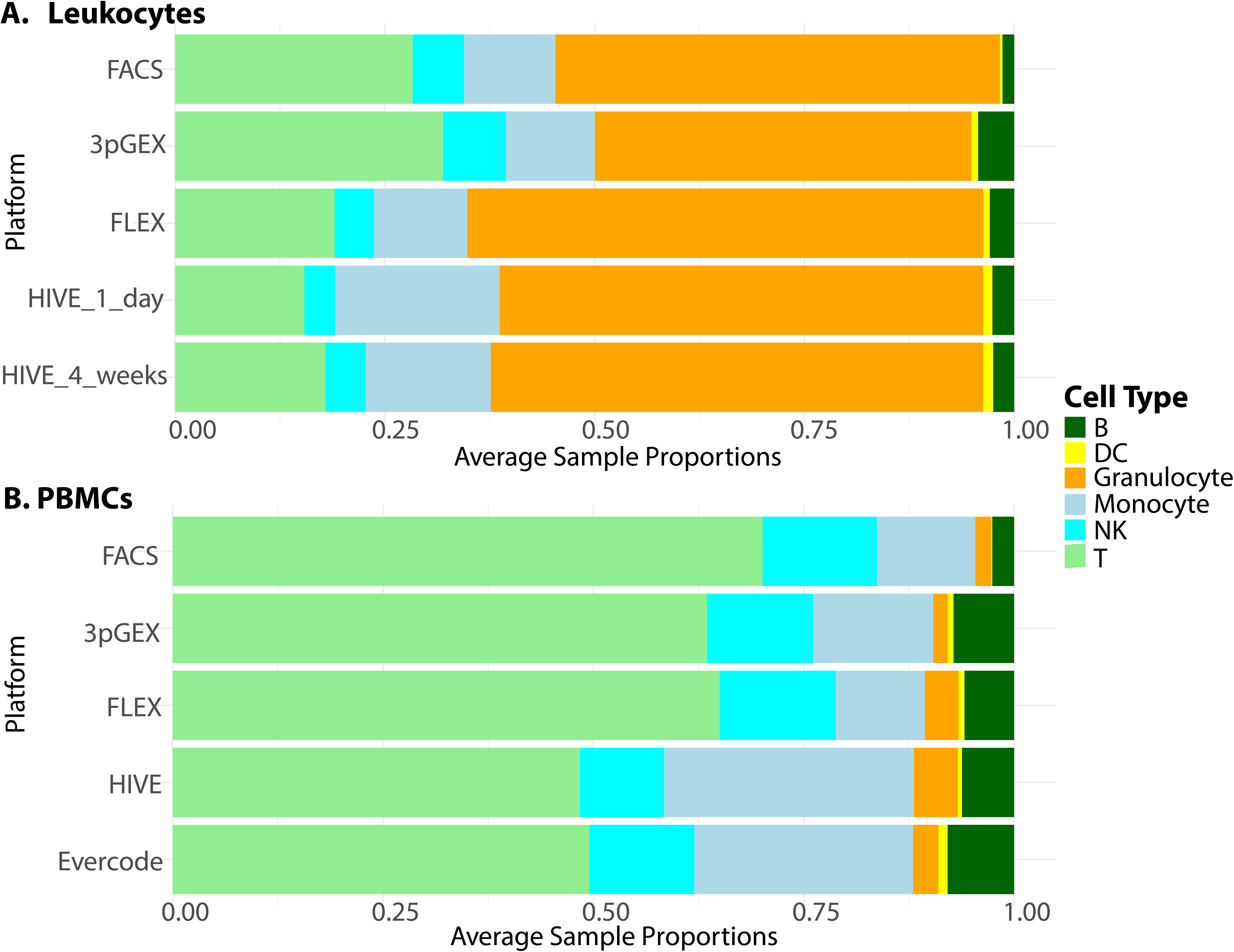
Comparison of cell-type proportions between scRNA-Seq and FACS. Cell type proportions in single cell RNA-Seq data aggregated by preparation compared to cell type proportions derived from FACS data for Leukocyte samples (A) and PBMCs (B). Although some variability exists in the proportion data, there is good agreement between the scRNA-Seq results and the FACS data. Cell type abbreviations are as follows: T=T-cell, B=B-cell, NK=Natural Killer cells, DC=Dendritic cells, The granulocyte proportions are consistently higher in the FLEX and HIVE platforms suggesting that these technologies more efficiently capture these fragile cell types.

To assess similarity of leukocyte expression profiles across different technologies, correlation analysis was performed on averaged data calculated using the ‘Seurat’ function *AggregateExpression* and the SCT assay. Importantly, genes not targeted by the 10x FLEX probeset were removed from this analysis. Correlation coefficients which were high across all technical replicates (minimum 0.93), sites (minimum 0.91) and technologies (minimum 0.91) with an overall minimum correlation coefficient of 0.75. Hierarchical clustering of correlation coefficients reveals strong clustering by technology, suggesting some level of platform specific expression (Fig. 5A). The HIVE and 10x data form two distinct clades, with the 1 day samples (10x 3pGEX) being distinct from the 28 day (FLEX). Within the two 28 day clades, the HIVE site samples and A and B replicates intermix with one another without discernible metadata associations suggesting that the site and technician variables have minimal contribution to the resulting expression data. However, within the 10x FLEX 28 day clade, site 1 and 4 samples are distinct from the site 2 and 3 samples and the A and B replicates within the site 1 and 4 samples are clustered together. This site distinction in the 10x FLEX data aligns with the QC parameter profiles (Fig. 3A) and may indicate that the 10x FLEX processing routine is more susceptible to technical variation from sample processing differences compared to the HIVE. Despite these differences, these data indicate each platform obtains leukocytes with highly similar expression profiles.

**Figure 5.**
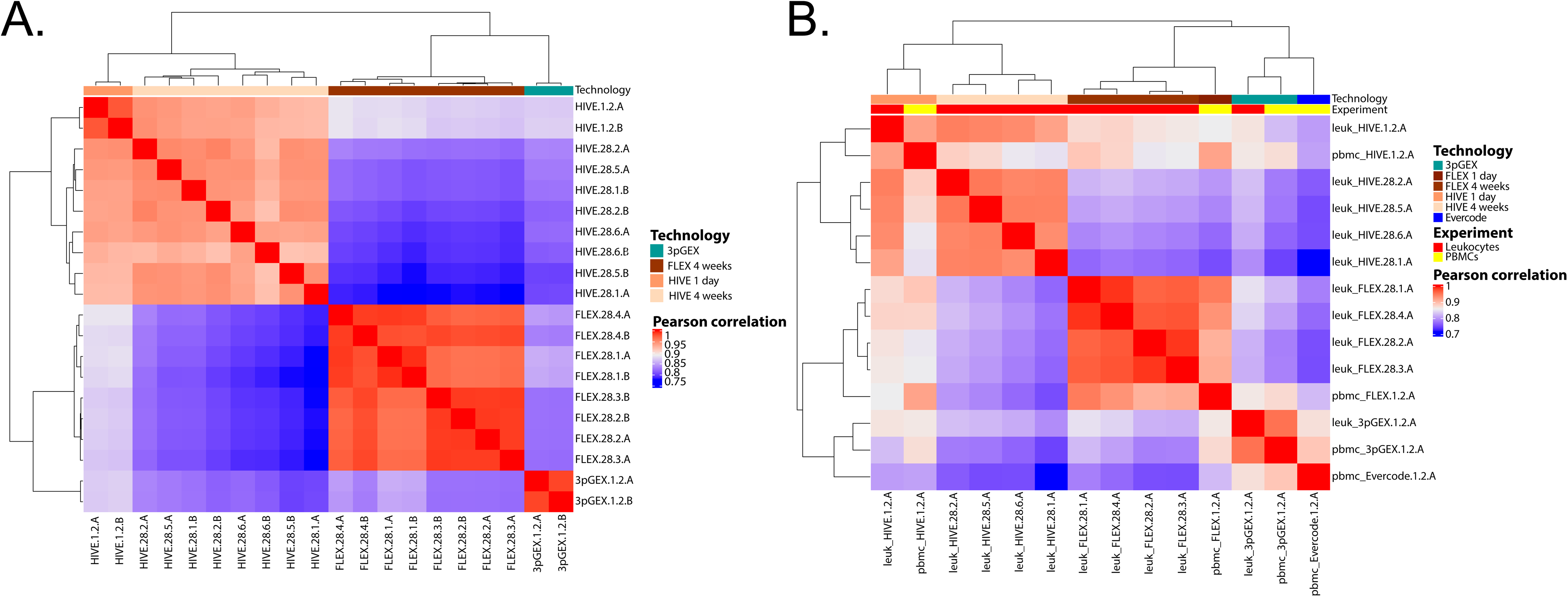
Correlation analysis of leukocyte data. Gene expression data for leukocyte samples was filtered to the subset of genes interrogated by the 10x FLEX platform and pairwise Pearson correlation coefficients were calculated (A). In the clustered heatmap, technology is indicated by the colored bar across the top of the heatmap. The HIVE and 10x samples group into two major clades. In the HIVE clade, the 1 day replicates are distinct from the 28 day replicates. Within the 28 day replicates, no association between the site-level A and B replicates is observed suggesting a high degree of reproducibility across sites. In the 10x clade, the FLEX samples are distinct from the 3pGEX samples. Within the FLEX clade, some association is observed between site-level A and B replicates for sites 1 and 4 while the A and B replicates are intermixed for the site 2 and 3 samples. Clustered heatmap of Pearson correlation coefficients for leukocyte A replicates and PBMC samples (B) shows technology-level clustering for various samples with leukocyte and PBMC samples clustered together for HIVE, day1, 10x FLEX and 10x 3pGEX technologies. The Parse Evercode PBMC sample is most highly correlated with the 10x 3pGEX samples. Only genes captured by FLEX probes were considered in the analysis. Technology and experiment are indicated by the colored bars across the top of the heatmap.

To investigate reproducibility of the fixation and storage process between replicates processed by different technicians at both the same site and across different sites, differential expression analyses between specific sample sets were performed (Fig. 6A). The violin plot in the top panel shows log 2 counts for each sample under consideration and the box plots in the bottom panel report the distribution of log2 fold changes for each A vs B comparison. The values for all tested genes are displayed and horizontal boxes at the 0 line indicate that more than 75% of the genes in the test have log fold changes near 0 and this is observed for all comparisons except for a single site (FLEX.28.4 A vs B). In this comparison, a subset of genes have fold changes > 0 indicated by the box above the 0 line and likely reflects differences in the counts highlighted by the violin plot in the top panel. Note that a small number of outliers were observed in each comparison. In total for all comparisons, there are a total of 12 outlier results that have fold changes outside the threshold of 1. These results come from 7 genes: HBB, IGHD, IGHG2, IGHG3, IGKC, IGLC3, and MT-CO1. It should be noted that reporting significant genes as defined by fold change and p-value thresholds does not work well for these analyses because most of the fold changes are small. Taken together, minimal differential expression between A and B replicates was observed indicating that all platforms are highly reproducible when processed in the same facility.

**Figure 6.**
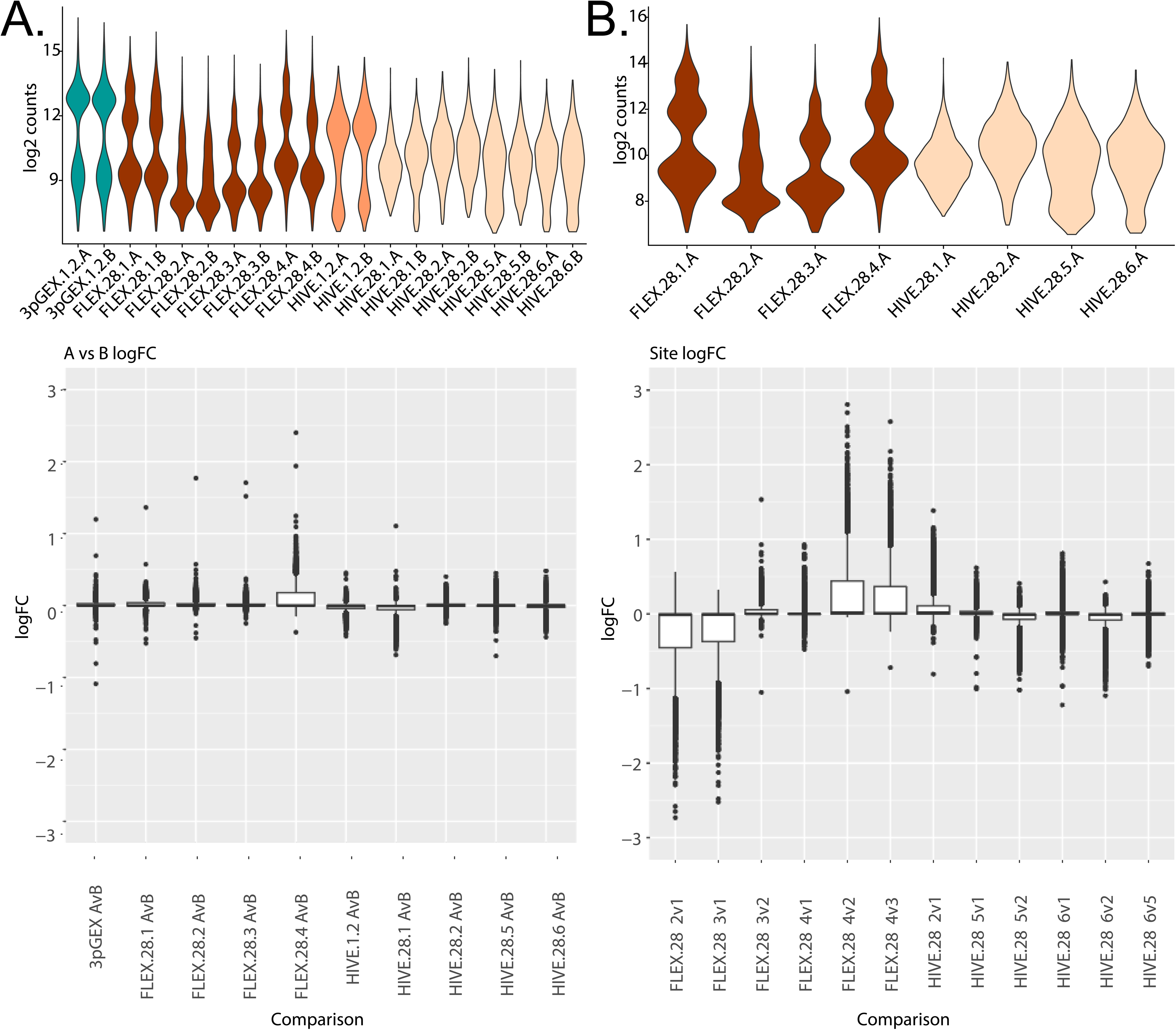
Differential expression analyses between site-level replicates and between sites. Differential expression analyses between the A and B replicates for each preparation and site (A) or between sites for each preparation (B). In both panels, violin plots of log2 counts per cell are provided to orient the comparisons being presented in the bottom panel box plots. The boxplots summarize the observed fold changes for each comparison. In many cases, the majority of the fold changes are near 0 indicated by a narrow box or line at 0 while comparisons with more substantial differential expression have a wider fold change distribution. For the A vs B comparison (A), the site 4 FLEX samples is the only comparison, only the FLEX.28.4 A vs B comparison has a substantial number of fold changes different than 0. In this case, non-0 fold changes are predominantly positive indicating higher values in A compared to B. This is consistent with the higher counts for FLEX.28.4A compared to FLEX.28.4.B. For comparisons between sites at 28 day storage (B) only A replicates were considered and each pairwise comparison within preparations was done. For FLEX, these comparisons were site 2 vs 1, 3 vs 1, 3 vs 2, 4 vs 1, 4 vs 2, and 4 vs 2. For HIVE, these comparisons were site 2 vs 1, 5 vs 1, 5 vs 2, 6 vs 1, 6 vs 2, and 6 vs 5. For the HIVE samples, comparison between sites does not produce many non-0 log fold changes. For FLEX however, any comparison between sites 1 or 4 with sites 2 or 3 produces genes with fold change. The log2 n_count plots show the count variability that underlies this result.

Differential expression analyses were also used to compare data generated at different sites (Fig. 6B). Given that the above correlation and differential expression data establish near-equivalence between technical replicates, site comparisons were performed using only the A replicates. The top panel shows the log2 counts across sites, with log2 fold-change values from all pairwise comparisons represented as boxplots in the bottom panel. For the FLEX samples, we observe elevated log2 UMI counts for the site 1 and site 4 samples, compared with those from sites 2 and 3. This distinction can also be seen when comparing the number of differentially expressed genes between these groups, with few differentially expressed genes detected when comparing sites 1 vs 4, and sites 2 vs 3, but a large number of differentially expressed genes comparing either site 1 or 4 with site 2 or 3. For the HIVE samples, none of the site comparisons produced substantial numbers of differentially expressed genes consistent with the correlation data presented above.

To further characterize library complexity of data generated across different platforms and sites and the differential expression result observed with site 4 in the 28 day FLEX samples, library complexity was modeled (Fig. 7A). 10X FLEX BAM files were downsampled and re-counted using a custom PySAM script that accurately reproduces Cellranger UMI calling logic. The resulting reads/UMI curve was fit using the Michaelis-Menten equation, where Vmax = Max # of detectable UMIs, and Km = Reads per Cell at 1/2 saturation. These Vmax values are plotted in the bar chart in Figure 7B and compared to the results for the 10x 3pGEX technology, filtering for genes included in the FLEX probe pool. Lower library complexity was observed in samples prepared at sites 2 and 3 compared to samples prepared at sites 1 and 4 l. This is supported by the barcode rank analysis, showing higher overall UMI counts in site 1 and 4 samples despite having grossly similar profiles (Fig. 7C).Substantial variability was also noted between A and B replicates from site 4 (Fig. 7D). Close examination of how FLEX samples were handled at each site revealed slight differences in the process that could contribute to the observed variation in UMI counts across sites. It was noted that sites 1 and 4 samples utilized fixed-angle rotors for centrifugation and appeared to have lower cell counts following probe hybridization, while sites 2 and 3 used swing-bucket rotors for these steps. These deviations in centrifugation do not appear to have an effect on the proportion of neutrophils from these samples which have lower RNA content and could account for this result, nor does it explain the variability between the A and B replicates only at site 4. Together these results indicate that the additional sample preparation steps required at each site for the FLEX assay can introduce variability and care should be taken to standardize these procedures as much as possible.

**Figure 7.**
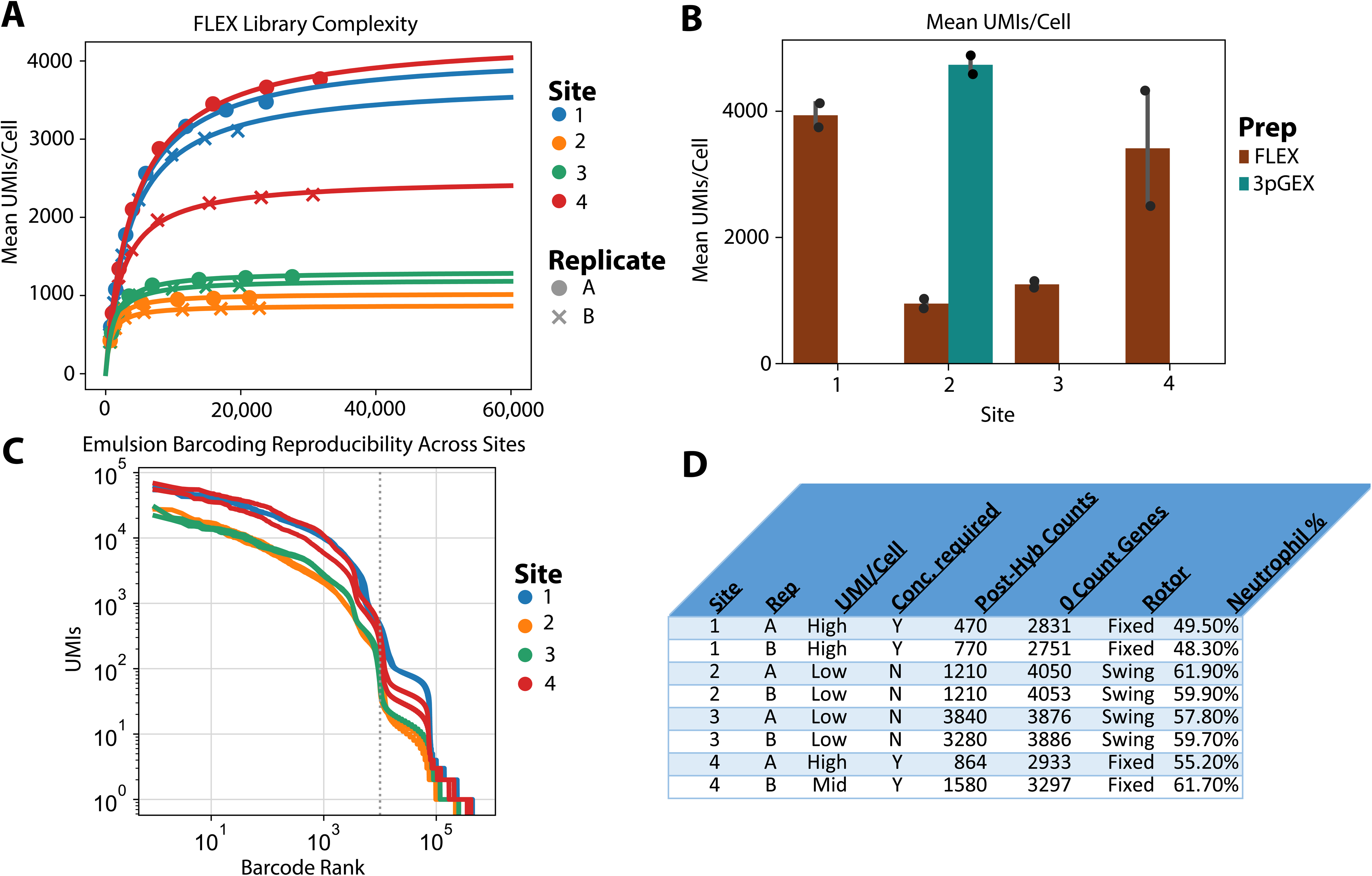
UMI analysis of 10x leukocyte samples. (A) The number of UMI per cell at different sequencing depths indicates variable library complexity in 10x FLEX leukocyte samples. Both site 1 samples and one of the site 4 samples have relatively high complexity, one site 4 replicate has intermediate complexity and all site 2 and site 3 samples have lower complexity. (B) The complexity of the site1 and 4 samples is similar to the 10x 3pGEX data. Bars show a modeled number of UMI per cell across sites and technologies, restricted only to genes that are captured by FLEX. (C) Plot of UMI compared to barcode rank indicates that approximately the same number of cells were loaded in each sample. (D) Experimental metadata from FLEX processing at each site.

Due to the incompatibility of RBC-depleted total leukocytes with the Parse Evercode v2 chemistry, an additional human PBMC sample was collected to allow assessment of all three technologies. Correlation analysis was performed (Fig. 5B). Samples clustered according to technology and to a lesser degree by sample type (i.e. leukocyte vs PBMC). This can be seen in the distinct clustering of FLEX and HIVE samples, as well as 3pGEX and Evercode samples, though the latter two technologies are most similar to one another. Interestingly, while HIVE 1d and 28d samples exhibit strong correlation coefficients (Pearson >0.9), they cluster separately from one another suggesting differences in expression profiles based on the length of storage prior to processing. To investigate this further, differential expression was performed between leukocyte HIVE day 1 and day 28 samples (Fig. S7) and no genes with log fold change > 1, were observed. These observations suggest that, while a high degree of consistency is observed, the choice of technology is not without impact and subtle differences in results can be detected.

In order to provide insight into the correlation analyses, the molecular biology underlying each method was considered in more detail using PBMC alignment data and the results are shown in the plots shown in Figure S8. The Evercode and HIVE alignments are roughly uniformly distributed along the transcript while, as expected, the 10x 3pGEX reads are concentrated at the 3’ end of the transcripts (Fig. S8A). The 10x FLEX alignments are probe-based, therefore average coverage distributions could not be calculated. Expression of a selected example gene (CD74, Fig. S8B) shows the expected coverage distribution for reads originating from each platform (Fig. S8C). This Integrated Genomics Viewer (IGV; Robinson et al. 2011; 2023) image of the CD74 locus with the PBMC BAM files loaded for display. 10x 3pGEX data appear at the 3’end of the gene, while 3 FLEX probes (red arrowheads, FLEX probe track) can be seen overlapping 3 internal exons. The centrally located probe spans an exon/intron junction resulting in a split appearance. The whole-transcript nature of the HIVE and Evercode reads is apparent from the alignments where coverage is highest over exons across the gene body.

Note, non-exonic coverage is observed in the 10x 3pGEX, HIVE and Evercode tracts. Examination of alignment in this region suggests that these reads may be caused by genomic regions rich in A or T bases (not shown) and is consistent with previous studies demonstrating internal priming of poly A/T tracts in nuclear pre-mRNA transcripts from technologies using oligo-dT based RT priming strategies (Svoboda et al. 2022).

## Discussion

Single-cell RNA sequencing has transformed our understanding of biology but has historically been limited to the analysis of high quality, fresh specimens processed at the site of collection. This requirement has hindered wide-spread adoption of this technique for clinical and other specimens where samples may be procured at sites without access to single cell instrumentation, or are collected over time and would benefit from additional data before committing to downstream processing. The availability of commercial solutions to preserve specimens prior to single cell analysis through cryopreservation and/or fixation addresses many of these issues, but systematic assessments of these approaches are lacking. Here we examine the ability of the 10x Genomics FLEX, HoneycombBio HIVE and Parse Evercode technologies to capture gene expression profiles from RBC-depleted total human leukocytes and PBMCs from a single individual, and compare the relative cell type proportions to 21-color flow cytometry data generated in parallel. While prior studies explored the differences between various scRNA-seq technologies, comparing metrics such as cost, performance, and sample compatibility (De Simone et al. 2024, Hatje et al. 2024, Xie et al. 2024, Hornung et al. 2023, Ding et al. 2020), ours is the first to assess the reproducibility of these methods in terms of both the sample preservation process, as well as the downstream capture and preparation of sequencing libraries across geographically separated core facilities.

In addition to the sample preparation challenges, components of downstream analyses were carried out in different core facilities. Because the deployment and management of consistent computing environments across sites can be difficult, we utilized a Docker image prepared by our team that contained all necessary software packages. This image was run on various high-performance compute environments using Singularity (Merkel 2014; Kurtzer, Sochat, and Bauer 2017). This approach enabled work to progress in different sites using identical software and it is an ideal for core facilities that wish to promote collaboration and reproducibility.

Both the 10x FLEX and Honeycomb HIVE kit performed well in preserving human leukocytes specimens, including retention of granulocyte populations, with relative proportions of cell types comparable to the “gold standard” flow cytometry dataset. While three of the four sites were able to generate sequencing libraries from total human leukocytes using the Parse technology, very few cells were captured and the data quality was poor (data not shown). That the PBMC samples were successful suggests that granulocytes may be problematic for this method. The findings from FLEX and HIVE are consistent with observations from Hatje et. al. and while they were able to generate libraries containing granulocytes, those populations are significantly underrepresented in their dataset. In contrast to Hatje et. al., we observe granulocytes in the 3pGEX dataset, though they are underrepresented compared to the other technologies. This result is likely due to the optimized sample collection procedure used for this study where cells were captured or preserved within 2hrs of the initial blood draw. Overall, the detection of granulocytes is an advantage of the FLEX and HIVE technologies compared with Parse or processing of fresh samples using 3pGEX. PBMCs were also examined using 10x 3pGEX, 10x FLEX, Honeycomb HIVE and Parse Evercode. All technologies produce high quality gene expression data that can be used to identify the expected cell types in proportions that are comparable to each other and concordant with flow cytometry measurements.

The ability to detect granulocytes has long been a challenge for single cell studies and even with their improved retention using preservation-based methods, computational solutions are needed to distinguish granulocyte-associated barcodes from those harboring ambient RNA or platelets, all of which have similar RNA content. Here we considered standard quality control parameters combined with analysis of cluster-specific markers to distinguish cell-associated droplets from acellular debris. This approach is effective at identifying cell-containing droplets and can be used to improve the recovery of RNA poor cell types independent of the platform used. Despite small exceptions, the cell type proportions we observe are similar in data from different platforms. This is consistent with previous reports that the 10x 3pGEX platform can capture granulocytes using specific sample preparation procedures and a modified analysis routine to include low RNA content cells (Salcher et al. 2024; Hatje et al. 2024).

Despite the high-level agreement between methods, detailed comparisons at the gene level reveal significant differences across technologies. Within the leukocyte dataset, correlation analysis between samples from 3pGEX, FLEX and HIVE demonstrate clustering by technology, with the stronger cross-site correlations observed for the 10x samples compared with HIVE. When including samples from the PBMC dataset, we again see strong clustering based on the technology used, rather than the sample type (i.e. leukocyte or PBMC). One possible explanation is the considerable difference in assay design across the technologies tested. We explored the distribution of read alignments and found that as expected, reads were concentrated in the 3’ region for samples prepared using 10x 3pGEX while the HIVE and Parse kits exhibit more uniform, full-length transcript coverage (Fig. S8). We also observed similar levels of mitochondrial abundance for both leukocytes and PBMCs with the exception of 10x FLEX which excludes many mitochondrial genes from the probe set (Fig. 2B). Despite these differences, broad concordance in cell type abundance, sample quality and total gene detection is consistent across technologies, enabling the successful integration of the datasets and comparison of QC metrics including gene and UMI counts per cell, which were similar across all technologies.

Our unique study design also allowed us to assess variation at the level of sample preservation, as well as across sites using the same technology. With the exception of the Site 4 FLEX replicates, we did not observe substantial differences in samples preserved by different technicians and analyzed at the same performance site (A vs B replicates, Fig. 5A.) We do note variability between samples analyzed by the same technology across sites. Specifically, 10x FLEX samples from sites 1 and 4 show higher UMI counts per cell than those processed at sites 2 and 3. While the precise reason for these differences is not known, we note that sites 2 and 3 used swing-bucket rotors for sample centrifugation, as opposed to fixed-angle rotors at sites 1 and 4, with the latter resulting in lower cell counts following probe hybridization. We hypothesized this cell loss may have resulted in a reduction in the neutrophil/granulocyte fraction and the capture of more high RNA content cells. However, no significant differences in neutrophil abundance were observed for these samples, though other populations may be affected (data not shown). This finding highlights the need to standardize downstream processing steps in particular for the FLEX method, where sample preservation is performed separately from single cell capture, which can introduce additional variability into the workflow.

The study presented here does not attempt to be a comprehensive quantitative assessment of the impact of preservation on scRNA-Seq data. Instead, we aimed to report on the experiences of multiple core facilities working with these technologies and to identify features of these workflows to be considered when deciding which to adopt for a given project. We find that in general, strong correlations exist between replicate samples processed at different sites using the same method, but that differences in expression exist between platforms. As a result, the method chosen should be based on the ability to accommodate the sample type and number of cells of interest, and logistics of sample collection and downstream processing. With PBMCs as input, all platforms produce high quality data as assessed by the ability to identify cell types in proportions consistent with the other technologies. For more challenging cell types such as total leukocytes, the 10x 3pGEX, 10x FLEX and HIVE platforms performed well based on the metric of cell type proportions. It is possible if users wish to minimize technical variability and all library preparations will be done at one location, 10x FLEX might be the best choice. If data will be produced at a variety of locations, Honeycomb HIVE may be the best option. It is also possible that the length of storage could contribute to variation in the data, as observed for the HIVE 1 day vs 28 day samples. This finding warrants further investigation across more timepoints and technologies to see if this effect is significant.

## Acknowledgments

The DSRG and GBIRG gratefully acknowledge the Association of Biomolecular Resource Facilities (ABRF) for funding our study, as well as the generous contributions provided by our commercial partners. This work would not have been possible without the reagents and technical expertise contributed by Parse Biosciences, Honeycomb Biotechnologies, 10x Genomics, and Illumina. CAW was partially supported by Cancer Center Support (core) Grant P30-CA14051 from the NCI to the Barbara K. Ostrom (1978) Bioinformatics and Computing Core Facility of the Swanson Biotechnology Center. MLM and SWP were supported by Delaware INBRE (NIH P20GM103446), Delaware CTR ACCEL (NIH U54GM104941), Delaware Biotechnology Institute, and the state of Delaware. FWK and OMW are supported by Cancer Center Support Grant (5P30CA023108) and NIGMS COBRE (P20GM130454) awards. The WashU GTAC@MGI is partially supported by NCI Cancer Center Support Grant P30CA91842 to the Siteman Cancer Center.

ABRF member cores that contributed to this work include; GSAF, University of Texas: RRID: SCR_021713, University of Delaware Bioinformatics Data Science Core Facility: RRID: SCR_017696, Dartmouth Genomics Shared Resource: RRID: SCR_021293, Center for Functional Genomics, UAlbany, Albany, NY. RRID: SCR_018262, Dartmouth Genomic Data Science Core: RRID: SCR_025383, Washington University School of Medicine Genome Technology Access Center Core Facility: RRID: SCR_001030, University of Wisconsin – Madison Biotechnology Center DNA Sequencing Facility: RRID: SCR_017759, ICBR Gene Expression & Genotyping at University of Florida: RRID: SCR_019145.

## Supplemental Materials

**Figure S1.**
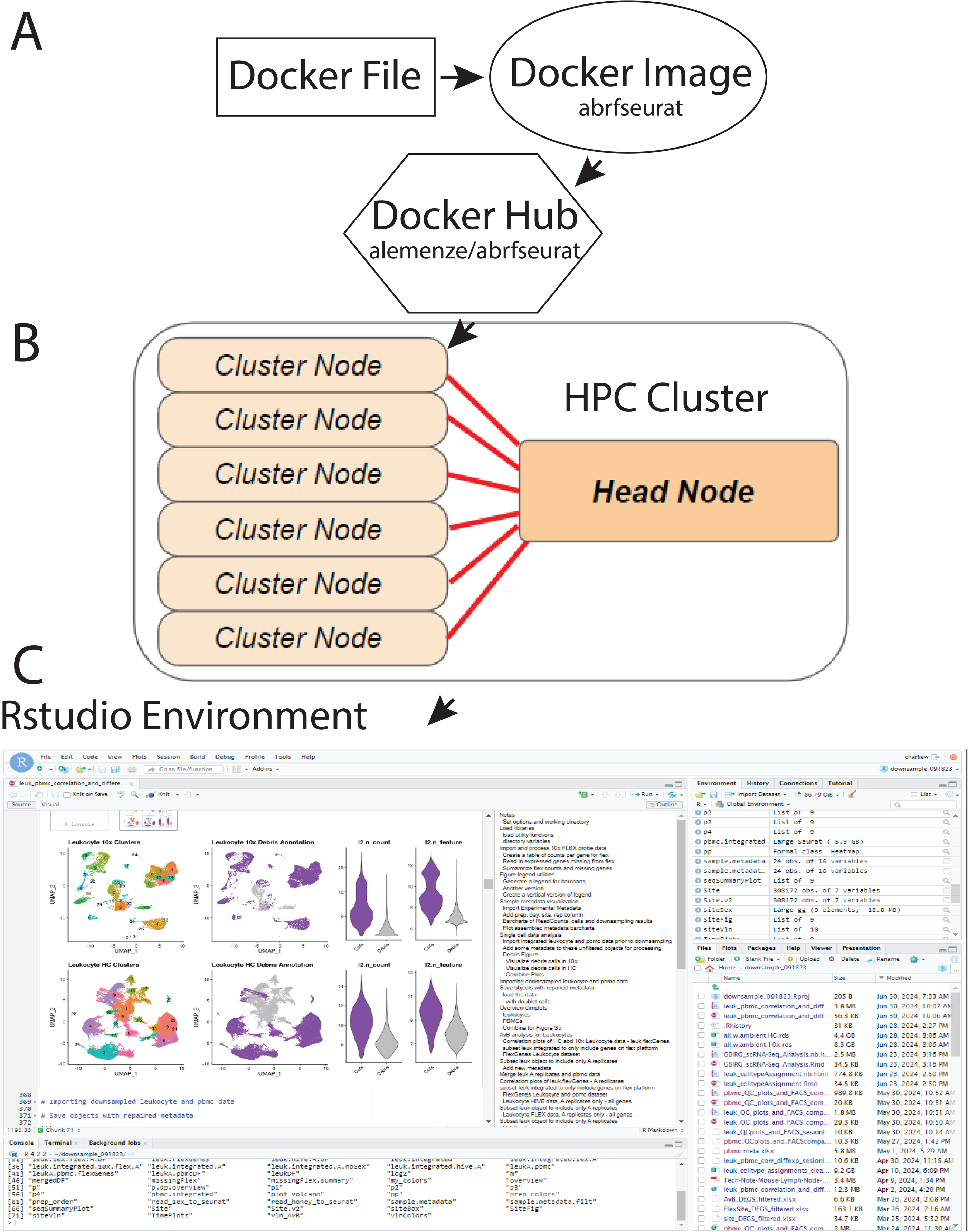
Container-based Rstudio analysis environment. Our study involved multiple collaborators at different institutions with various computer infrastructures and required a computational solution to provide a consistent analysis environment. A) A Docker Rstudio image containing all required software packages was prepared and deposited in Docker Hub (docker://alemenze/abrfseurat). B) This container can be executed using Docker, Singularity, or Apptainer in any computational environment. Given the scale of the data in this study, the most common environment used was a Linux-based high performance computer (HPC) cluster. C) After launching the R environment, contributors can interact with the software from the command-line or by using the browser-based Rstudio interface.

**Figure S2.**
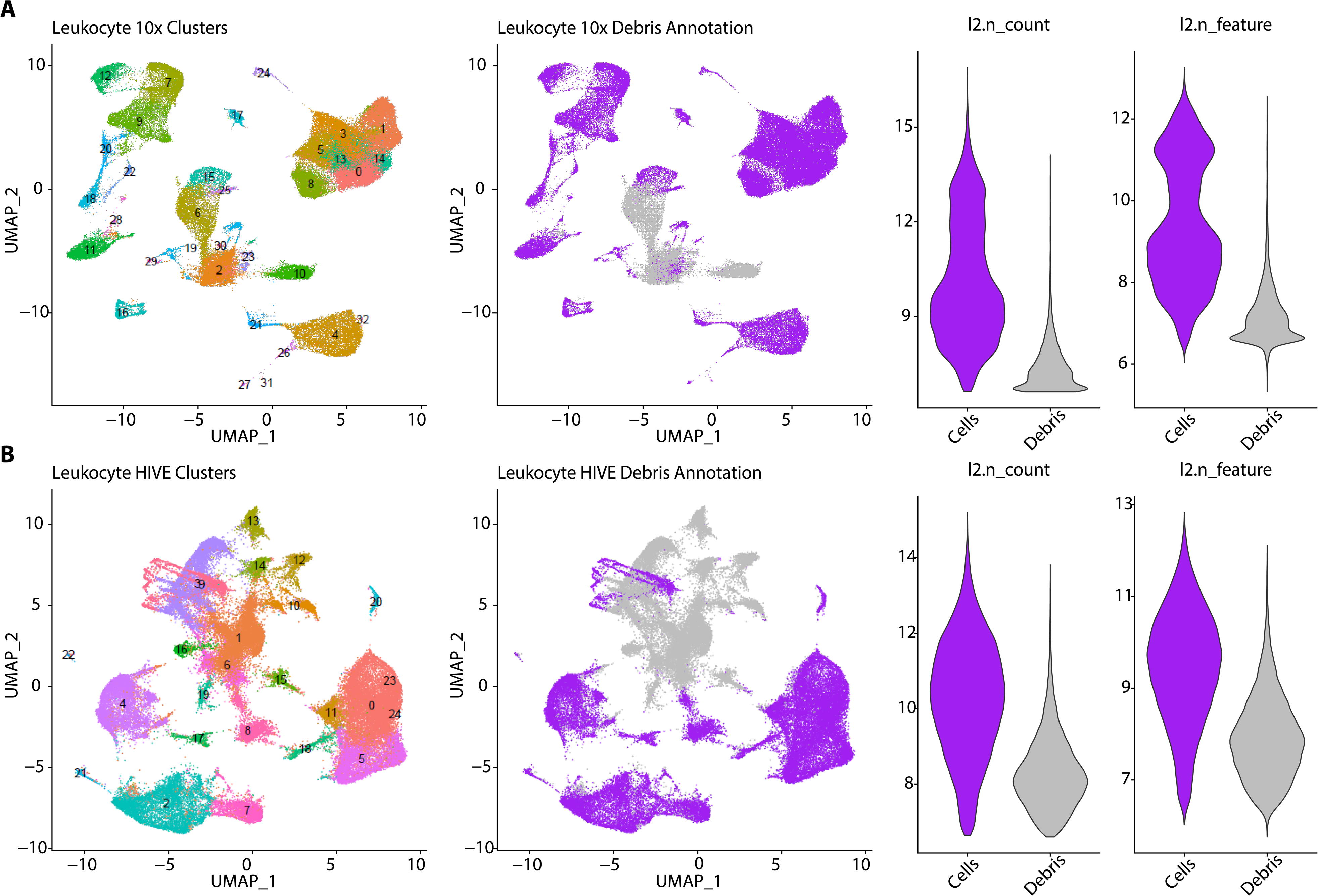
Cluster-level debris exclusion. Low quality leukocyte clusters with ambiguous cell identity were excluded from analysis prior to downsampling. A) Leukocytes from 10x 3pGEX or 10x FLEX and B) Leukocytes prepared with HIVE technology. Left panels are cluster assignments used to evaluate real vs debris clusters. Middle panels are the same UMAPs but colored according to real cell flag (purple) or debris (grey). Right panels are violin plots of l2.n_count and l2.n_feature statistics stratified by debris assignment using the same color scheme as the adjacent UMAPs plots. Debris clusters are generally in the center of the UMAP and have poor quality control statistics compared to real cells.

**Figure S3.**
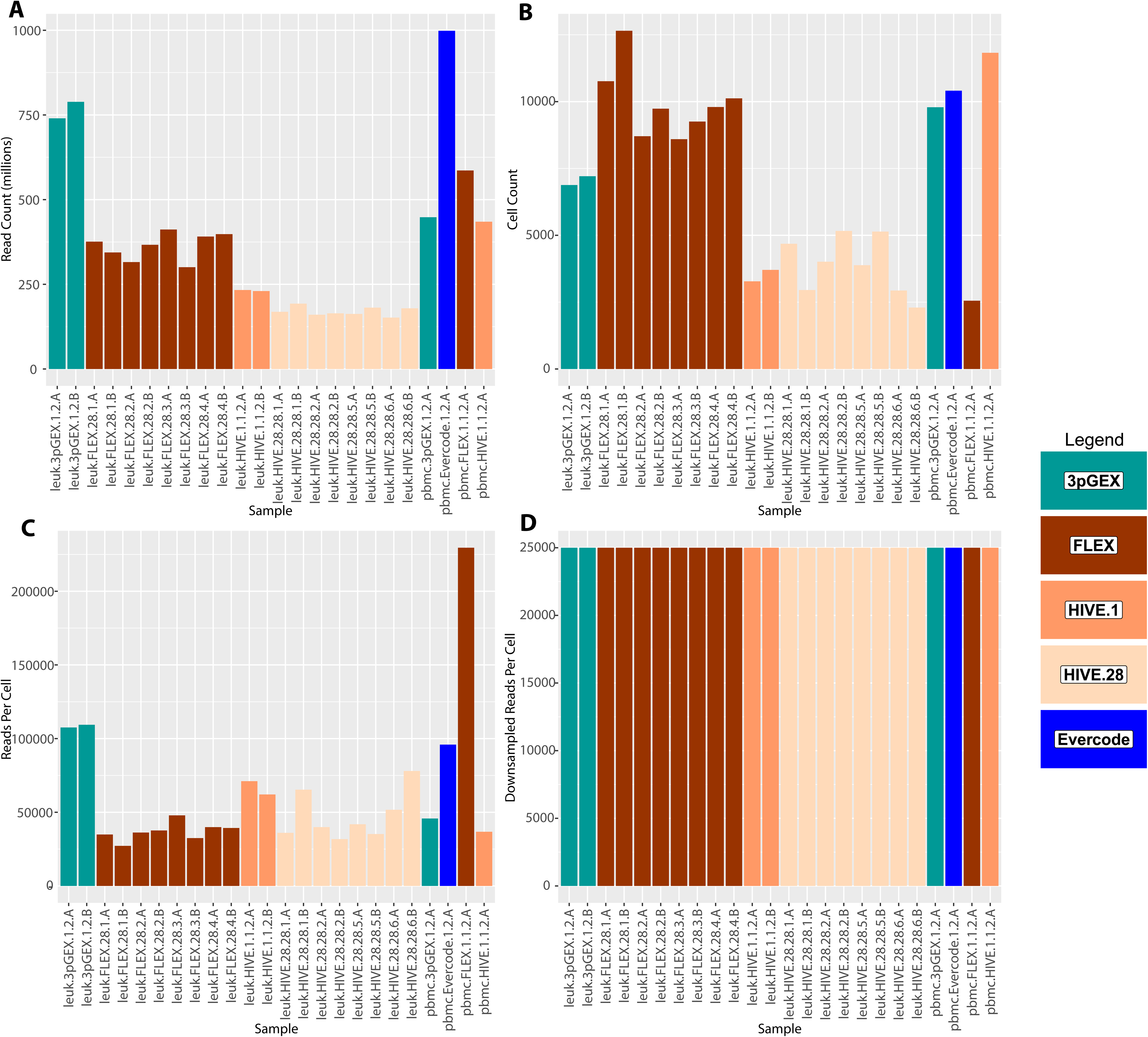
Sequencing summary. Bar charts of different sequencing parameters for the 24 samples considered in this study. In each panel, colors are according to technology and storage parameters. The leukocytes are the 20 bars to the left, PBMCs are the 4 bars to the right. A) Overall depth of sequencing in millions of reads. B) Cell counts per sample after debris exclusion. C) Sequencing depth in reads per cell prior to downsampling input FASTQ files. D) Sequencing depth in reads per cell after downsampling FASTQ files to 25,000 reads per cell. These downsampled FASTQ files were then used as input in all analyses.

**Figure S4.**
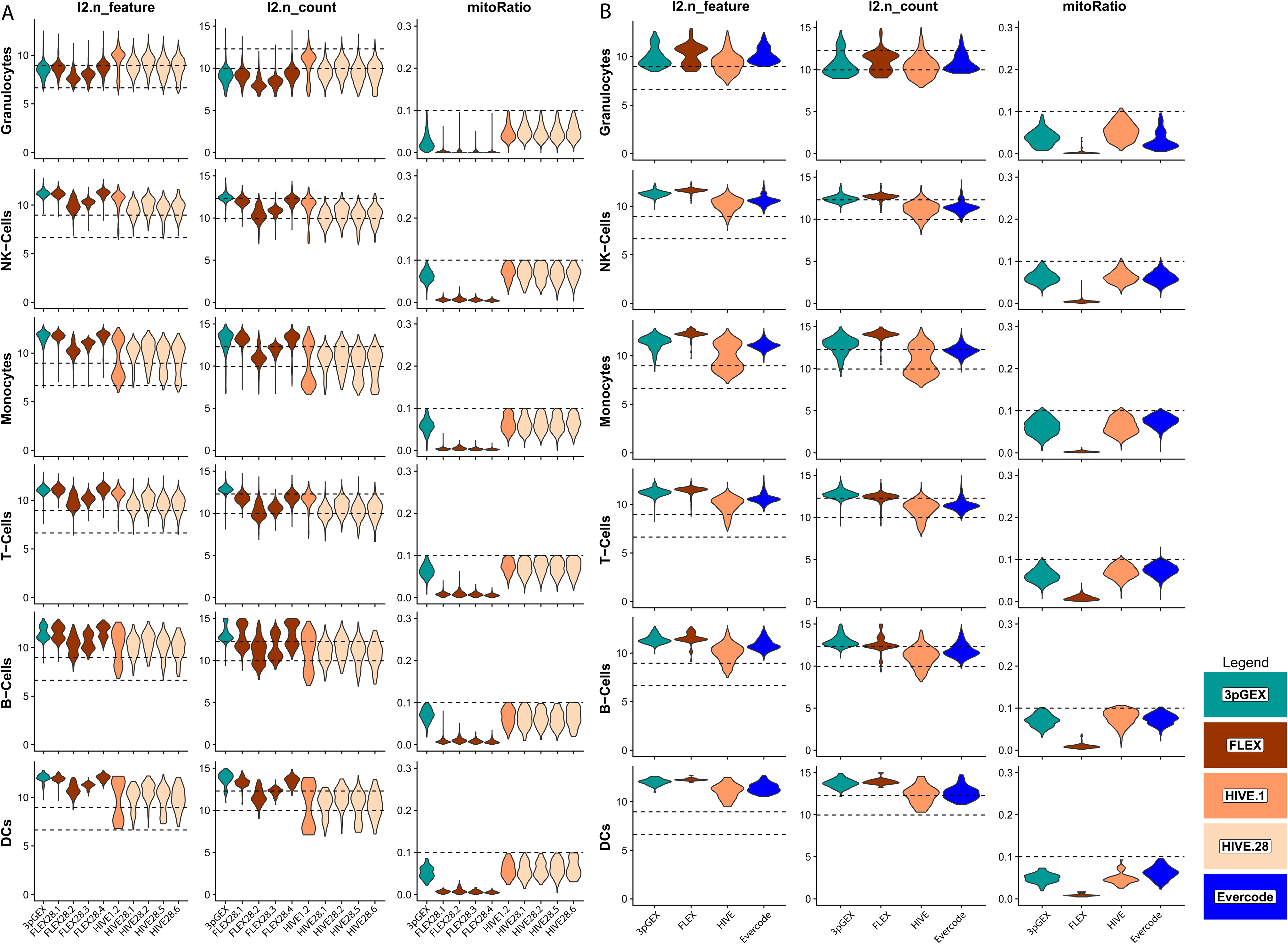
Quality control statistics for Leukocyte and PBMC samples stratified by general cell type. Violin plots presenting the log2 transformed gene count (l2.n_feature), read counts (l2.n_counts) and the percent mitochondrial reads (mitoRatio) for filtered and downsampled leukocyte (A) and PBMC (B) samples. The leukocyte site-level replicates have been merged for this presentation and the values for each general cell type are plotted separately. Mitochondrial read percentages are relatively low in the 10x FLEX samples because probes interrogating these genes are excluded from the platform. Violins are colored according to technology. Dashed reference lines are provide at 6.6 (100) and 9 (500) for gene counts in l2.n_feature panels, 10 (1000) and 12.3 (5000) for read counts in l2.n_count panels and 0.1 (10%) for percent mitochondrial reads in mitoRatio panels.

**Figure S5.**
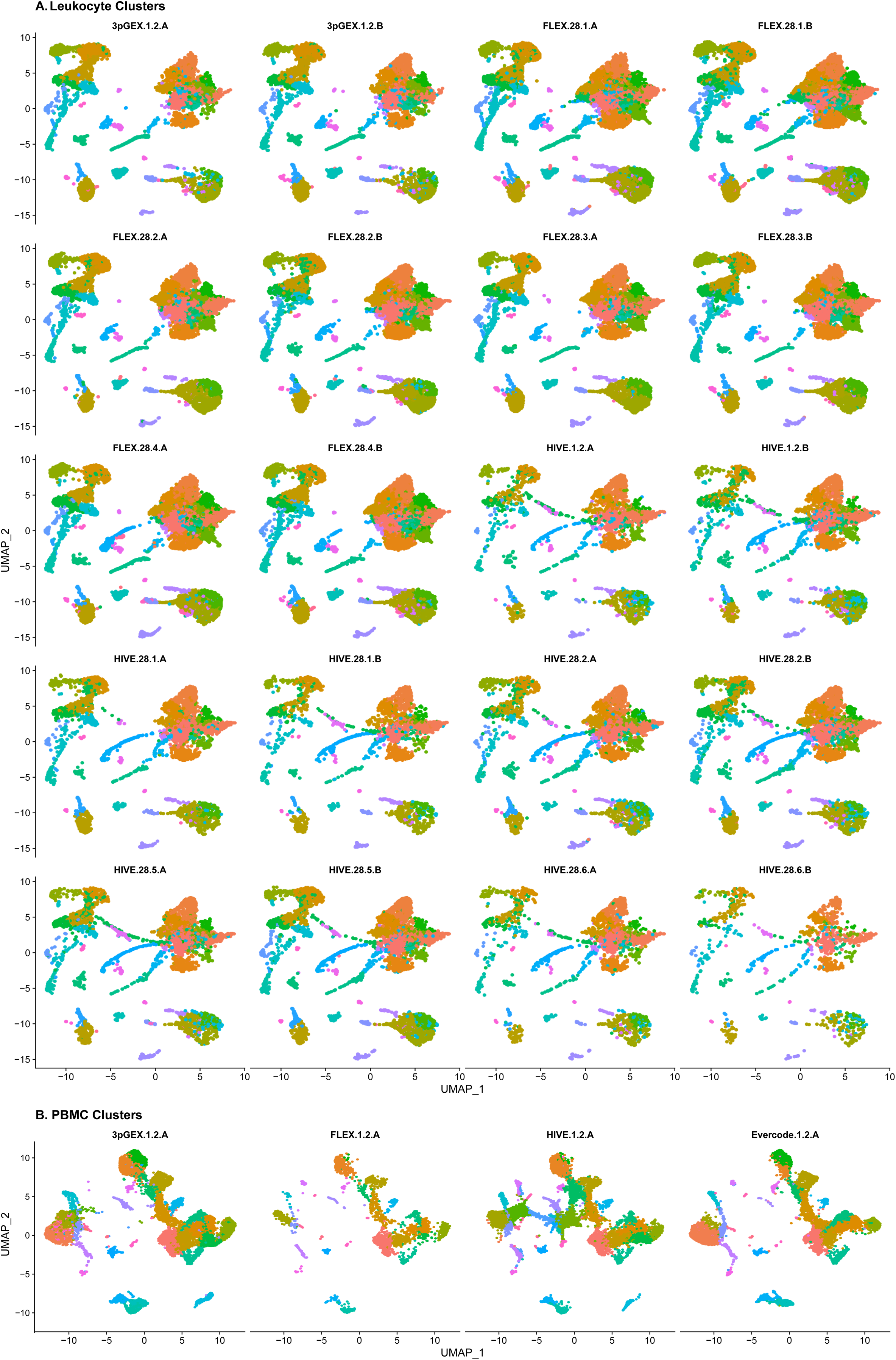
Post-integration UMAP plots of each sample. Following integration of leukocyte (A) and PBMC (B) data, UMAP plots show similar distributions of cells in these two dimensional representations indicating successful integration of data from different technologies. UMAP plots are colored according to cluster.

**Figure S6.**
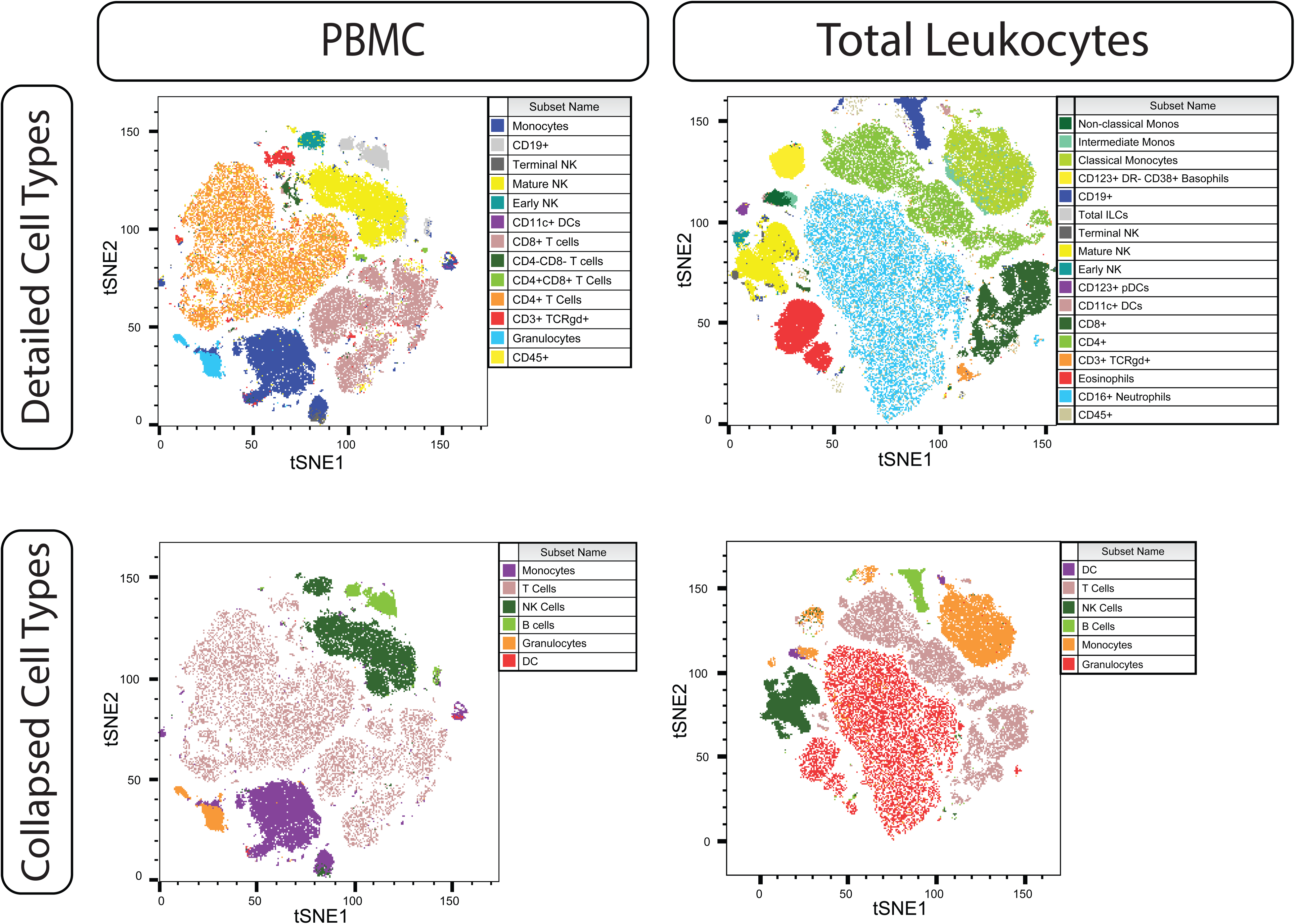
FACS tSNE plot. The CD45+ leukocyte population as characterized by flow cytometry using a 21-color panel of surface marker antibodies (See Table S1).

**Figure S7.**
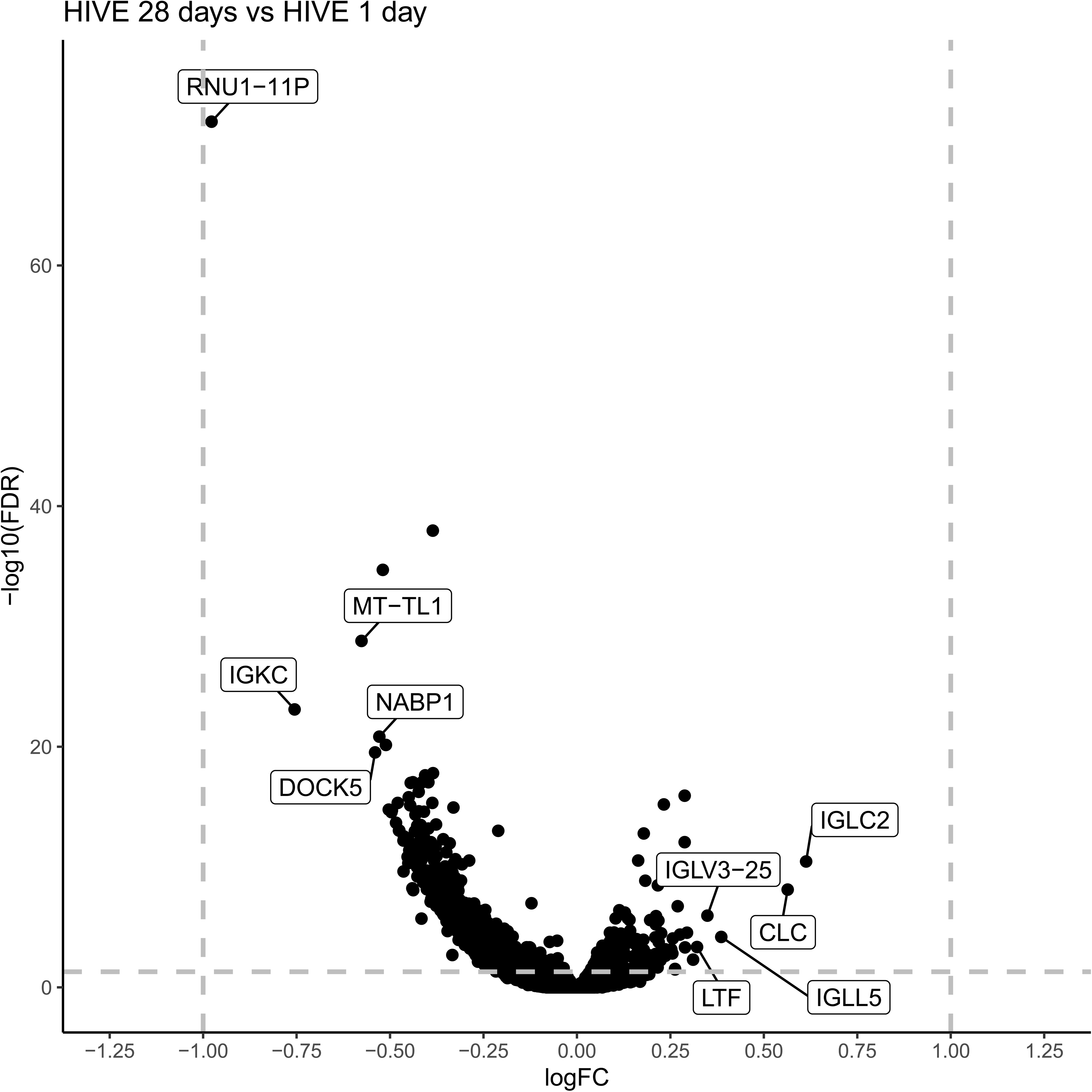
Differential expression analysis of HIVE 4 weeks vs HIVE 1 day leukocyte samples. Volcano plot of differential expression results between the HIVE 28 day storage A replicates and the HIVE 1 day A replicate. Genes with the largest magnitude positive and negative log2 fold changes are labeled. No genes exceed the abs(log2 Fold Change) > 1, FDR < 0.05 threshold for differential expression.

**Table S1:** Flow Cytometry Antibodies.

| SPECIFICITY | FLUOROCHROME | CLONE | CATALOG # | VENDOR |
| --- | --- | --- | --- | --- |
| CD45RA | BUV395 | 5H9 | 740315 | BD |
| CD16 | BUV496 | 3G8 | 564653 | BD |
| CD56 | BUV737 | NCAM16.2 | 564447 | BD |
| CD8 | BUV805 | SK1 | 612889 | BD |
| CD197 (CCR7) | BV421 | G043H7 | 353208 | BioLegend |
| CD123 | Super Bright 436 | 6H6 | 62-1239-42 | ThermoFisher |
| CD11c | eFluor 450 | 3.9 | 48-0116-42 | ThermoFisher |
| IgD | BV480 | IA6-2 | 566138 | BD |
| CD3 | BV510 | SK7 | 344828 | BioLegend |
| CD28 | BV650 | CD28.2 | 302946 | BioLegend |
| CD14 | Spark Blue 550 | 63D3 | 367148 | BioLegend |
| CD45 | PerCP | HI30 | 368506 | BioLegend |
| TCR$\gamma\delta$ | PerCP-eFluor 710 | B1.1 | 46-9959-42 | ThermoFisher |
| Siglec-8 | PE | 7C9 | 347104 | BioLegend |
| CD4 | cFluor YG584 | SK3 | R7-20041-100T | Cytek612889 |
| CD25 | PE-CF594 | M-A251 | 562403 | BD |
| HLA-DR | AF647 | L243 | 307622 | BioLegend |
| CD19 | Spark NIR 685 | HIB19 | 302270 | BioLegend |
| CD127 | R718 | HIL-7R-M21 | 352753 | BD |
| CD27 | APC-H7 | M-T271 | 560222 | BD |
| CD38 | APC-Fire810 | HIT2 | 356643 | BioLegend |

**Figure S8.**
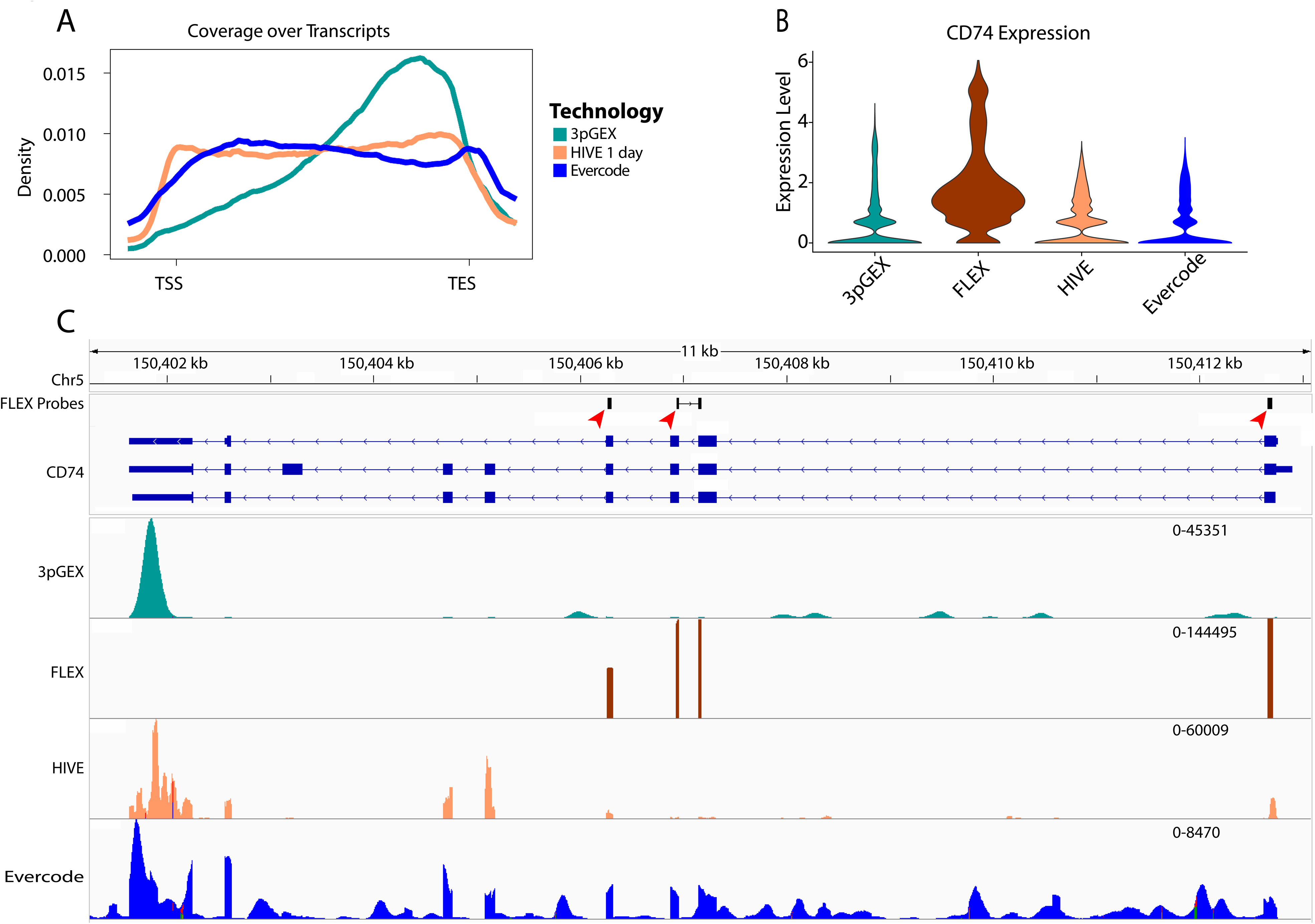
Molecular biology differences between technologies. Examination of the different molecular biology features of the technologies tested in the PBMC samples in this study. A) Transcript metagene plot showing depth of coverage from transcript start (TSS) to transcript end (TES) for all expressed transcripts from each technology. The 10x 3pGEX align near the TES while HIVE and Evercode data are more uniformly distributed across the entire transcript length. The 10x FLEX data is not included in this plot because these data are derived from probes and are not compatible with whole transcript coverage plots. B) Expression levels of the example immune gene CD74 in each technology. Normalized and log10-transformed data from the data slot of the SCT assay are being presented. C) Sequence alignment data visualized using IGV in the CD74 example locus from each technology. The 10x 3p DGE reads mostly localize to the 3’ end of the negative-stranded CD74 gene. The 10x FLEX alignments are restricted to the location of the FLEX probes (Red Arrowheads). Note that the middle probe is split between two exons and this results in a corresponding pair of read stacks. The HIVE and Evercode alignments are observed across the length of the transcript, predominantly over exons. Non-exonic alignments are often near genomic regions that are rich in A or T suggesting spurious genomic priming (data not shown).

